# Spontaneous HFO Sequences Reveal Propagation Pathways for Precise Delineation of Epileptogenic Networks

**DOI:** 10.1101/2024.05.02.592202

**Authors:** Zhengxiang Cai, Xiyuan Jiang, Anto Bagić, Gregory A. Worrell, Mark Richardson, Bin He

## Abstract

Epilepsy, a neurological disorder affecting millions worldwide, poses great challenges in precisely delineating the epileptogenic zone – the brain region generating seizures – for effective treatment. High-frequency oscillations (HFOs) are emerging as promising biomarkers; however, the clinical utility is hindered by the difficulties in distinguishing pathological HFOs from non-epileptiform activities at single electrode and single patient resolution and understanding their dynamic role in epileptic networks. Here, we introduce an HFO-sequencing approach to analyze spontaneous HFOs traversing cortical regions in 40 drug-resistant epilepsy patients. This data-driven method automatically detected over 8.9 million HFOs, pinpointing pathological HFO- networks, and unveiled intricate millisecond-scale spatiotemporal dynamics, stability, and functional connectivity of HFOs in prolonged intracranial EEG recordings. These HFO sequences demonstrated a significant improvement in localization of epileptic tissue, with an 818.47% increase in concordance with seizure-onset zone (mean error: 2.92 mm), compared to conventional benchmarks. They also accurately predicted seizure outcomes for 90% AUC based on pre-surgical information using generalized linear models. Importantly, this mapping remained reliable even with short recordings (mean standard deviation: 3.23 mm for 30-minute segments). Furthermore, HFO sequences exhibited distinct yet highly repetitive spatiotemporal patterns, characterized by pronounced synchrony and predominant inward information flow from periphery towards areas involved in propagation, suggesting a crucial role for excitation-inhibition balance in HFO initiation and progression. Together, these findings shed light on the intricate organization of epileptic network and highlight the potential of HFO-sequencing as a translational tool for improved diagnosis, surgical targeting, and ultimately, better outcomes for vulnerable patients with drug-resistant epilepsy.

**One Sentence Summary:** Pathological fast brain oscillations travel like traffic along varied routes, outlining recurrently visited neural sites emerging as critical hotspots in epilepsy network.

## INTRODUCTION

Epilepsy, a prevalent neurological disorder, affects around 70 million people globally (*1*). Accurate localization of the epileptogenic zone (EZ, the brain area and connections required for seizure generation) is crucial for successful surgical outcomes (*2*). Current clinical practice relies on intracranial electroencephalography (iEEG) recording using electrocorticography (ECoG) from the subdural surface of the brain and/or stereo-EEG (sEEG) from the brain parenchyma to identify pathological and physiological activities essential for localizing the epileptic brain during pre-surgical planning (*3*). This process, typically spanning 5-14 days, involves reducing anti-seizure medication to record the habitual seizures (*4*). However, the extended periods of invasiveness and discomfort, along with various risks associated with seizure provocation and chronic electrode implantation, poses substantial drawbacks (*5, 6*). Moreover, the rapid spread of seizure signals can obscure the precise onset of epileptic activity (*7, 8*). Despite careful patient selection, about 40- 50% of patients do not experience favorable outcomes after surgery (*3, 6, 9*). This suggests a discrepancy between the identified seizure onset zone (SOZ) and the actual EZ, highlighting the critical need for accurate and reliable biomarkers to improve surgical success rates.

High-frequency oscillations (HFOs) in electrophysiological recordings have emerged as a promising biomarker for the EZ (*8, 10*) with close relation to the SOZ (*11, 12*). Characterized as spontaneous oscillations above 80 Hz lasting for at least four cycles (*13*), HFOs are correlated with ictogenesis (*14*), and their removal from brain areas has been linked to improved surgical outcomes (*15–17*), potentially offering a degree of precision surpassing traditional markers, e.g., irritative zone (IZ) or SOZ (*18, 19*). However, significant challenges remain at the single patient and single electrode level resolution for mapping epileptogenic brain (*20*), particularly in differentiating clinically relevant HFOs from physiological (*21–23*) or artifact-related oscillations (*13, 24, 25*) due to their overlapping frequency ranges (*26, 27*). Moreover, this issue is compounded when HFOs are identified solely by their occurrence rates, as physiological HFOs may be mistaken for pathological ones, reducing their specificity (*28, 29*) and even leading to conflicting findings (*30–32*). Additionally, the common practice of manual HFOs marking, particularly in short data segments, is labor-intensive and susceptible to subjective biases (*33, 34*). While HFOs detectors have been proposed (*35–37*), these approaches are mainly semi-automated and require expert review to achieve optimal performance, highlighting the need for more refined and automated methods in HFO analysis.

Recent research into various forms of HFOs, including stereotypical HFOs (*38, 39*), spike ripples (*36, 40–42*), and onset HFOs (*11, 43, 44*), as well as physio/pathological distinctions (*23, 45*), has brought attention to their unique characteristics and morphologies in relation to epilepsy. These findings suggest that the complexities of HFOs offer new angles to comprehend the epileptic brain, extending beyond occurrences. Yet, a critical limitation of prior studies is the reliance on short data segments, which may overlook the broader spectrum of HFO-behavior and variability over time. This has contributed to ongoing debates regarding the significance of HFOs in comprehensive, extended recordings (*31*).

In contemporary clinical practice, there is a pressing need for reliable methods to identify pathological HFOs and accurately delineate epileptogenic network at the individual patient level. Crucially, a deep understanding of HFOs, including their origins and intricate spatiotemporal dynamics, is vital. This necessitates an examination of HFO stability in extended recordings and detailed investigation into their spatiotemporal organization at a fine-grained, millisecond scale. Leveraging on previous studies and ours (*40, 43, 44*), we observed spontaneous HFOs traversing cortical regions within short timeframes, forming sequenced pathways in iEEG recordings. This study aims to bridge these gaps and provide a nuanced understanding of HFOs in the context of epilepsy.

In the present study, we introduce a data-driven approach to systematically sequence and analyze propagational HFOs over extensive iEEG recordings from patients with drug-resistant focal epilepsy (Fig. 1A,B). Our objective is to precisely identify the origins, pathways, and the cortical regions implicated in these HFO sequences and to scrutinize their propagation patterns (Fig. 1C,D). We hypothesize that spontaneous HFO sequences are a reliable marker of the EZ, offering new perspectives on the initiation and progression of HFOs and on the broader epileptic network. This study includes a thorough examination of HFO sequences to assess their efficacy and reliability as biomarker for localizing epileptic tissues and their spatiotemporal dynamics and functional connectivity. By doing so, we seek to address and clarify ongoing debates regarding HFOs and devise methods in accurately pinpointing the EZ, thereby enhancing the precision of epilepsy surgery and improving patient management and treatment outcomes.

**Figure 1.**
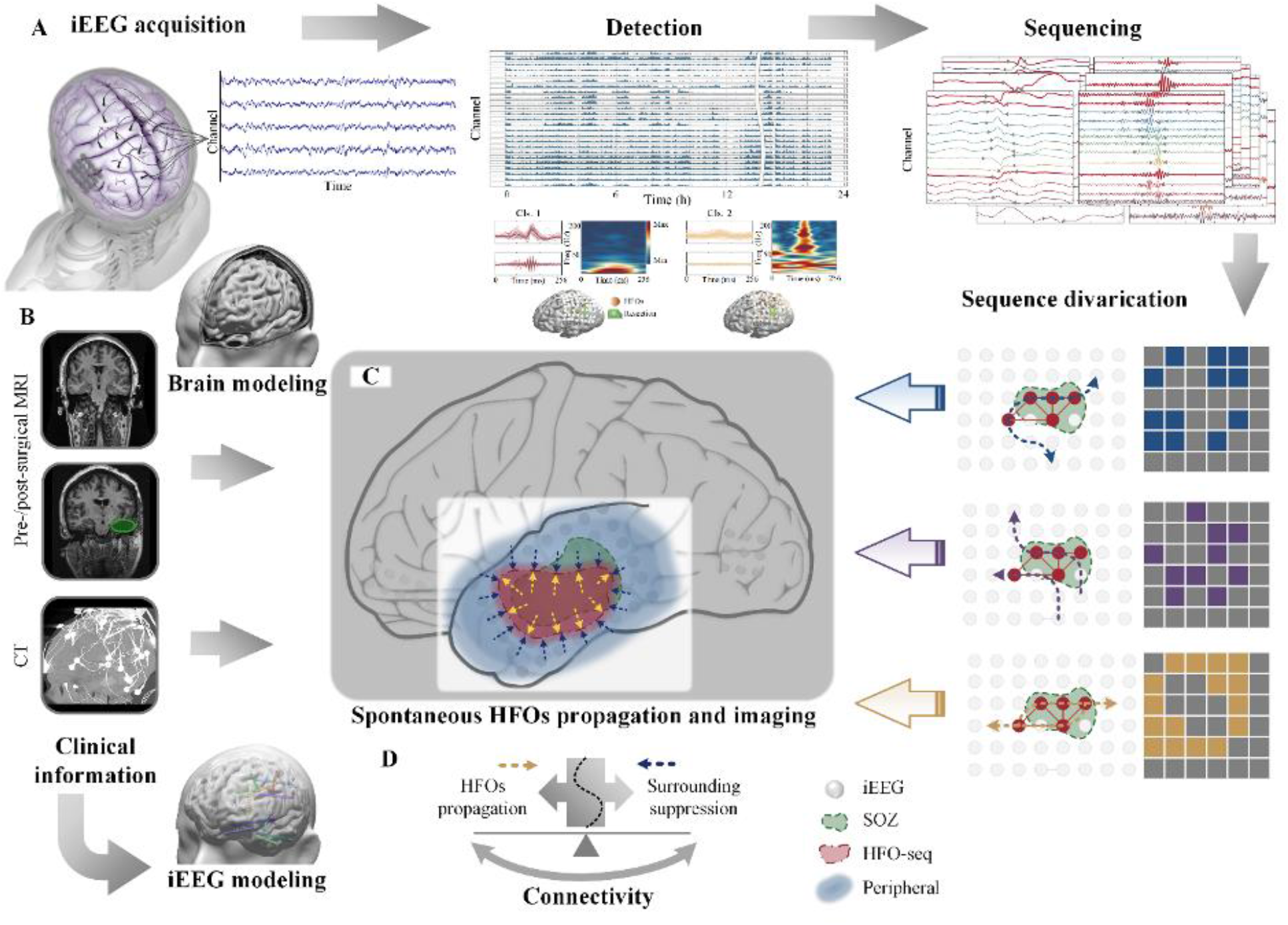
Conceptual diagram of data analysis and hypothesis. **(A)** iEEG data analysis workflow. Long-term iEEG data were collected and HFOs were detected and sequenced to identify the propagation of HFOs, which consists of a series of HFOs being captured across multiple channels spanning a tight time window in a relatively local cortical area. **(B)** Clinical evidence and modeling. Pre-surgical MRI was used to build a realistic head model for individual patients, CT images were co-registered with the pre-surgical MRI and used to localize the iEEG implantation from which the SOZ was defined by the clinicians, and post-surgical MRI was for modeling the surgical resection zone (green). **(C)** Mapping of spontaneous HFO sequences. The main hypothesis is that sequenced HFOs are concurrent propagating activities highly associated with the pathological zones of epilepsy, by which the underlying tissue epileptogenicity can be delineated. Multiple spatiotemporal divarication may exist, while the highly recruited core region of HFO sequences provides precise and stable mapping of the epileptogenic networks. **(D)** Connectivity antagonism between the HFOs and surrounding areas. The information flow between the HFO-zone and outside is associated with the propagation of the HFOs, and hence reflects the underlying pathology of the epileptic network.

## RESULTS

### Propagation of spontaneous HFOs in human subjects

During our clinical investigation, HFOs were identified to manifest simultaneously across multiple electrodes, forming sequenced HFO patterns within a localized spatial and temporal span in long-term iEEG recordings. To study the behavior and properties of such sequenced HFOs, which consist of a series of HFOs captured across multiple channels within a short time frame, we conducted extensive spontaneous (interictal) epileptic activity recordings from iEEG in a cohort of 40 drug-resistant focal epilepsy patients. These recordings were part of presurgical monitoring across two clinical centers. For each patient, an average recording length of 23.40 ± 2.51 *hours* (*mean* ± *SD*, *n* = 40) was analyzed, with equal attention to daytime (11.69 ± 1.10 *hours*, *mean* ± *SD*) and nighttime periods (11.70 ± 1.64 *hours*). Detailed descriptions of patient information and data collection are included in Materials and Methods.

In total, 101,185.59 channel hours of iEEG data from 40 patients were analyzed, averaging 106.95 ± 37.62 channels of iEEG per patient (*mean* ± *SD*, *n* = 40). To conduct HFOs detection and sequencing, we developed an automated and integrated approach based on established methods with modifications (see Materials and Methods) (*40, 46, 47*). This approach employed high sensitivity detection and unsupervised learning techniques to capture spontaneous interictal HFOs, characterized by at least four cycles of oscillations distinct from the background, showing an isolated power increase in the high frequency band (>80 Hz), in line with standard clinical study criteria (*10, 13*). The detected HFOs were then aggregated and categorized by their temporal and spatial instances to form HFO sequences, identifying those concurrent events co-occurring across multiple electrodes within a short timeframe, representing spatiotemporal co-activation or propagation of HFO activities.

In this process, a total of 8,936,866 putative HFO events were detected across all patients, with a comparable number of activities during the day and the night (*p* = 0.71, *n* = 40, two-sided Wilcoxon signed-rank test). The vast dataset generated through these recordings presented a wide spectrum of activities – pathological, physiological, and spurious (*13, 21, 24*) – due to the shared frequency bands and complex morphological variations in extended recordings (*25–27*). In line with previous research (*31*), the detected HFO events spanned extensive areas or multiple regions, with certain channel groups showing recurrent patterns of simultaneous HFO detection (Fig. S1A).

Importantly, the clustering process of detected events identified apparent artifacts, characterized by high amplitude and sharp activities, which were subsequently excluded before further analysis (Fig. S1B). Echoing prior findings (*38, 40, 41, 48*), we also noted the presence of HFOs with various repetitive patterns within these clusters (Fig. S1B).

Next, to track the propagation of HFOs, we aggregated the initial detections and sorted the HFO series that were aligned spatiotemporally and occurred in sequences spanning multiple electrodes within a short timeframe (Fig. 2; Materials and Methods). Across all patients, 273,697 HFO sequences (HFO-seq) were identified, with an average 7.73 ± 1.27 events per sequence (*mean* ± *SD*, *n* = 40). Notably, a higher proportion of sequences occurred during the night compared to that of all detected HFOs (aHFOs: 47 ± 13.74%, HFO-seq: 52.09 ± 15.84%, *mean* ± *SD*, *n* = 40; *p* = 0.0028, two-sided Wilcoxon signed-rank test). This agrees with prior studies of increased pathological HFOs during sleep (*49, 50*), less affected by motion-related artifacts (*13*). Each identified HFO sequence typically covered a local extent of about 55.74 ± 23.89 *mm* with an average propagation time of 67.58 ± 22.58 *ms* (*mean* ± *SD*, *n* = 40; Fig. 2). Besides, diverse spatiotemporal patterns were observed in HFO-seq (Fig. 2), with repetitive instances involving similar electrode pairs within individual patients, suggesting consistent propagation pathways across HFO networks (Fig. S2). Overall, our proposed approach proved efficacy and generalizability in detecting and sequencing HFOs across the study cohort, enabling the automatic identification of varied sequences in prolonged iEEG recordings.

**Figure 2.**
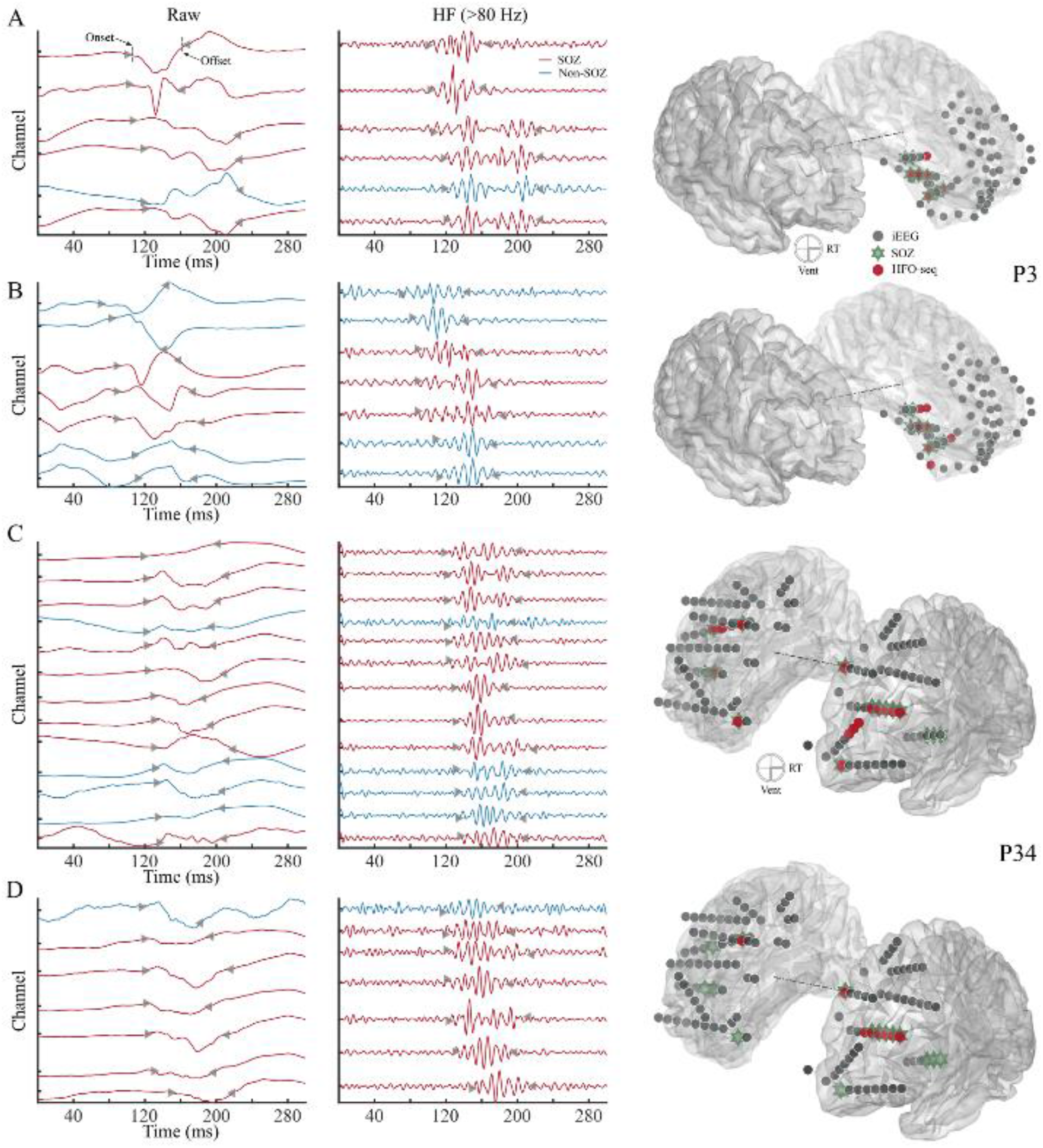
Examples of identified HFO sequences. (A,B) Two HFO sequences from P3. **(C,D)** Two HFO sequences from P34. The examples showcase distinct HFOs morphology between sequences. The left panel of each row shows the raw signal data, and the middle panel displays the signals after high-pass filtering above 80 Hz. The red line indicates SOZ channels, and blue as non-SOZ channels. Onset and offset of events are denoted by left and right-pointing triangles, respectively. The rightmost panel for each row visualizes the spatial positioning of iEEG channels (dark) on the cortical surface—channels participating in the sequences are highlighted in red, and the SOZ is marked with a green hexagon.

### HFO-seq elevates precision of occurrence rates for SOZ association

First, we evaluated the detected HFOs and HFO-seq using the conventional occurrence rate, which quantifies the count of activities per minute (*51*). The main hypothesis is whether the identified HFO-seq is a refined biomarker compared to the total occurring HFOs in the clinical settings. To address this, occurrence rate was calculated for all detected HFOs (aHFOs) and HFO sequences (HFO-seq) in 10-minute segments across the prolonged recording for each patient with a benchmark association to the SOZ, the brain area where seizures originate, determined by the clinical experts. We observed that the aHFOs generally showed variable clusters in different spatial locations over time, while the HFO-seq displayed more distinct and localized activation patterns, predominantly highlighting the activities relevant to SOZ (Fig. 3A). This suggests a more precise SOZ association with HFO-seq compared to general HFOs.

**Figure 3.**
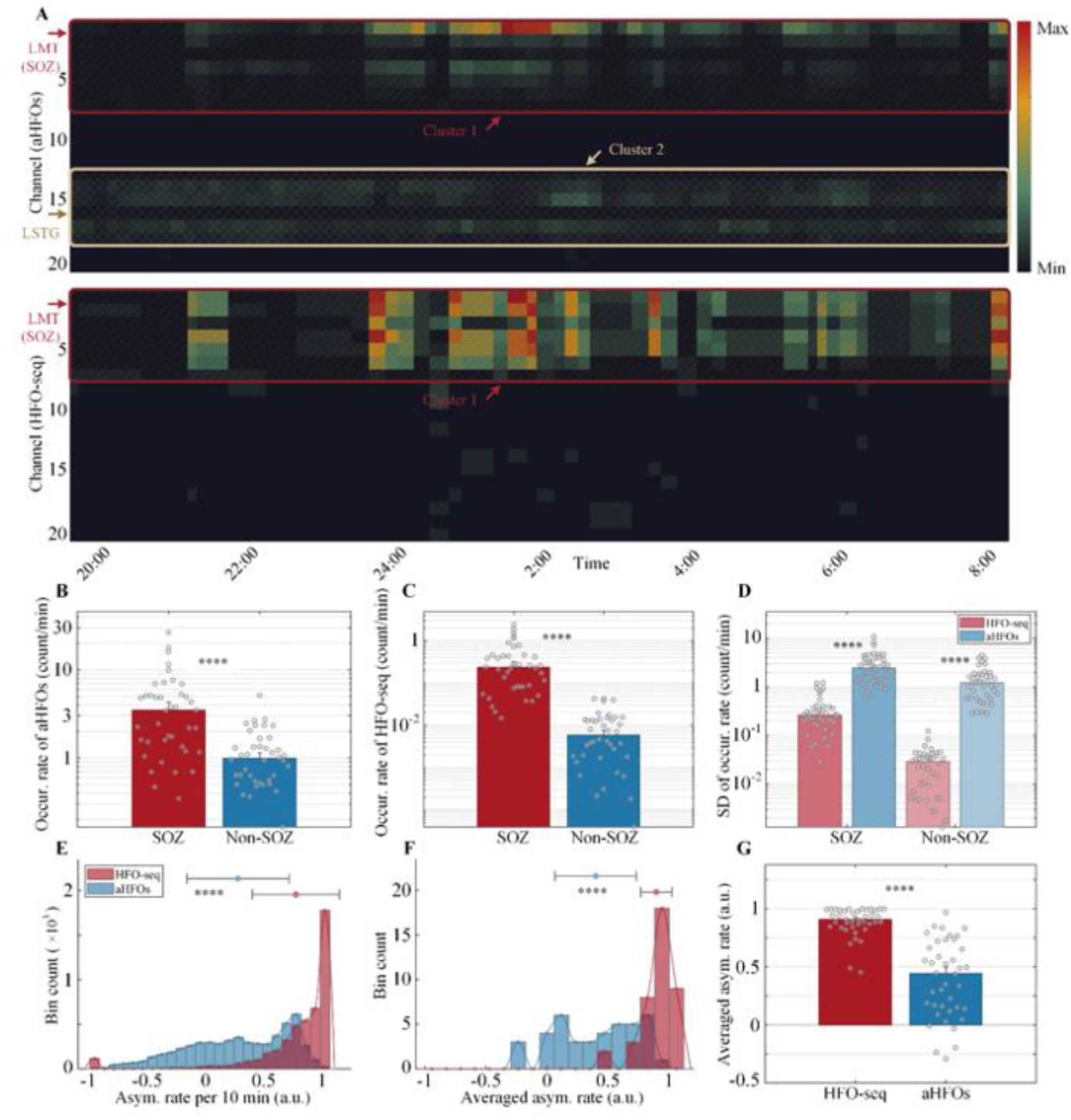
Occurrence and asymmetry rate of detected HFOs and identified HFO sequences. (A) Raster plot of HFO occurrence rates for all detected HFOs (aHFOs, upper panel) and HFO sequences (HFO-seq, lower panel). The rows represent recording channels, and columns denote time indices. The occurrence rate is color-coded, with the scale representing the number of events per 10 minutes. Notably, the SOZ channels are displayed at the top. Distinct clustering patterns are observed in the aHFOs, with one major cluster associated with the SOZ (hippocampal area) and another emanating from the posterior temporal region, while HFO-seq predominantly occupies the hippocampal site, demonstrating clear spatial foci. **(B,C)** Occurrence rates of aHFOs and HFO-seq, respecitvely, within SOZ versus non-SOZ. Note the seperation of SOZ versus non-SOZ between aHFOs and HFO-seq. **(D)** Standard deviation of the occurrence rates calculated from every 10-minute segment throughout a full day. Note the order of magnitude of the standard deviation in both measures under logarithmic scale. **(E,F)** Histogram of asymmetry rates of aHFOs and HFO- seq measured acroess every 10-minute segments and entire recording duration in all patients (*n* = 40). **(G)** Averaged asymmetry rate between HFO-seq and aHFOs across the patient cohort. ∗∗∗∗ *p* < 0.0001.

In our patient cohort, the occurrence rate of aHFOs within the SOZ was significantly higher than outside it (SOZ: 4.75 ± 5.31 HFOs/min, *mean* ± *SD*, non-SOZ: 1.23 ± 0.95 HFOs/min; *n* = 40, *p* < 10^−6^, two-sided Wilcoxon signed rank test; Fig. 3B), aligning with existing literature suggesting close relation of HFOs to the SOZ (*52*). A similar while stronger trend was observed for HFO-seq, where rates inside the SOZ were higher compared to outside (SOZ: 0.43 ± 0.54, non-SOZ: 0.01 ± 0.01; *mean* ± *SD*; *n* = 40; *p* < 10^−7^, two-sided Wilcoxon signed rank test; Fig. 3C). Besides, the standard deviation of occurrence rates for HFO-seq was notably lower than for aHFOs in both SOZ and non-SOZ areas (*n* = 40, both *p* < 10^−7^, two-sided Wilcoxon signed rank test; Fig. 3D), indicating a significantly low variability in HFO-seq.

Moreover, we assessed the asymmetry index of the occurrence rates for both aHFOs and HFO-seq as another established metric (*31*), which quantifies the correlation between the activity rate and the SOZ (Fig. 4E,F; Materials and Methods). Both aHFOs and HFO-seq showed positive asymmetry, indicating notable association with the SOZ. However, the distribution for aHFOs was more spread out compared to HFO-seq, whose distribution was tighter towards one. Significantly, HFO-seq exhibited a markedly higher asymmetry rate towards the SOZ than aHFOs (aHFOs: 0.39 ± 0.33, HFO-seq: 0.89 ± 0.13; *mean* ± *SD*; *n* = 40; *p* < 10^−7^, two-sided Wilcoxon signed rank test; Fig. 4G). This result reinforces our earlier findings, highlighting the greater pathological relevance and stability of HFO-seq as a refined biomarker.

**Figure 4.**
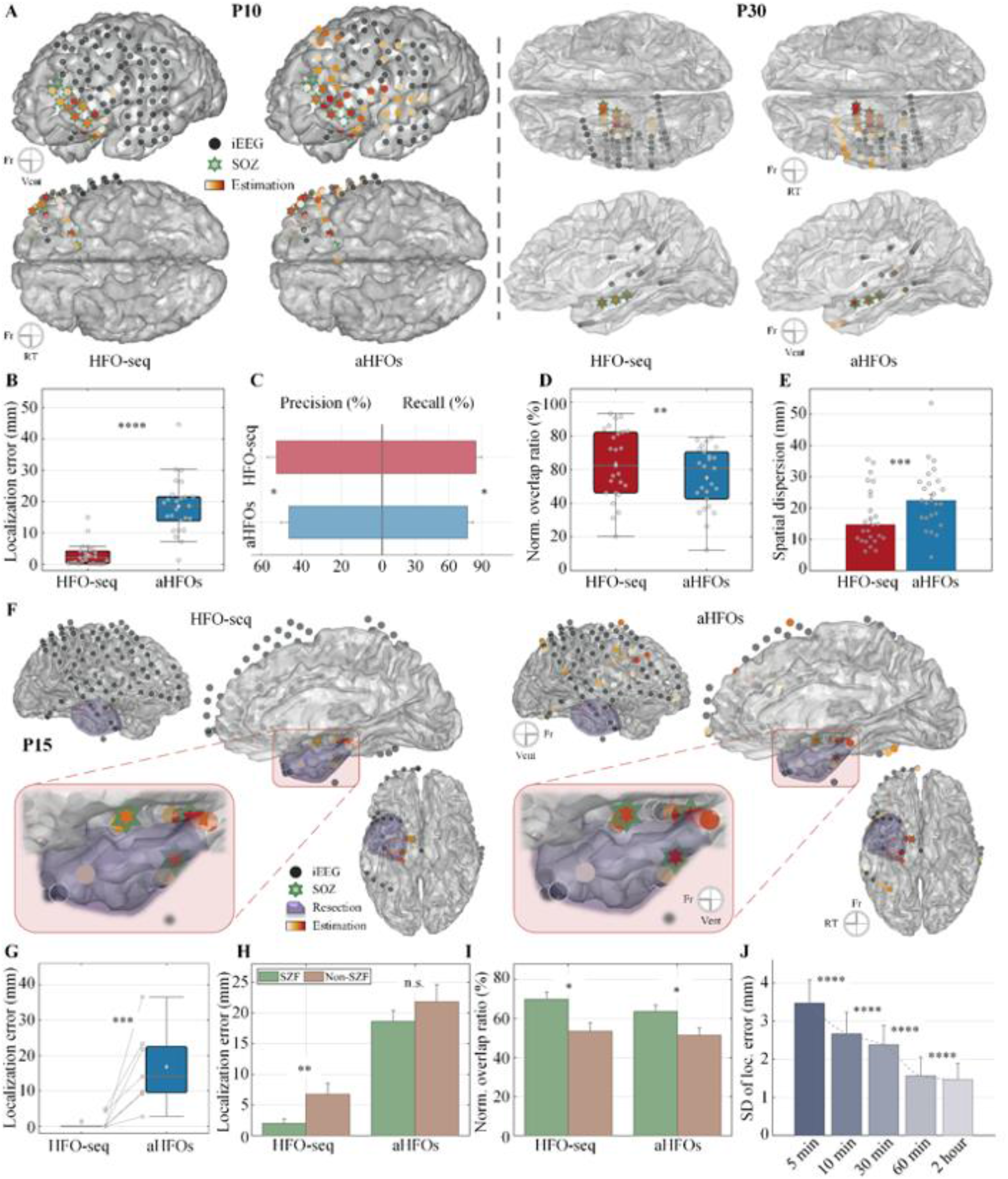
Localization of SOZ and resection zone using HFO-seq and aHFOs. (A) Mapping of HFO sequences (HFO-seq) and all HFOs (aHFOs) to localize the SOZ (green stars). The iEEG electrodes are shown in dark, with one case for ECoG (left panel) and one for sEEG (right panel). The estimations are color coded from light (minimal activity) to warm (maximal activity). **(B)** Localization error of HFO-seq and aHFOs in relation to the SOZ. The overlap between the estimated HFO locations and the SOZ is evaluated through: **(C)** Precision and recall, **(D)** Normalized overlap ratio, and **(E)** Spatial dispersion. **(F)** Mapping of HFO-seq and aHFOs in relation to the resection zone (purple). **(G)** Localization error of HFO- seq and aHFOs to the resection. **(H,I)** Comparison of imaging performance to the SOZ associated with different seizure outcomes, seizure-free (SZF) and non-seizure-free (non-SZF). **(J)** Standard deviation of localization across different segment lengths using HFO-seq mapping. Note the remarkable decrease from 5 to 10 minutes and from 30 minutes to 1 hour. Fr: frontal, Vent: ventral, RT: right temporal. ∗ *p* < 0.05,∗ ∗ *p* < 0.01,∗∗∗ *p* < 0.001,∗∗∗∗ *p* < 0.0001.

### Localization of epileptic tissues through HFOs mapping

To investigate whether the sequenced HFOs (HFO-seq) provide accurate localization of epileptogenic tissues compared to the conventional HFOs (aHFOs), we quantified their cortical activation distribution and juxtaposed to the clinically presumed EZ, including the SOZ for 25 patients and the resection volume for seven patients with seizure-free outcomes. In these cases, the EZ approximation was based on the SOZ region and/or resection areas, considering the successful surgical outcomes (*3*). We employed multiple metrics for evaluation, including localization error (LE), normalized overlap ratio (NOR), and spatial dispersion (SD) (*40, 53–58*), to assess the performance of both HFO-seq and aHFOs in identifying the EZ. The methodology and metrics are elaborated in Materials and Methods.

Overall, the estimations of both modalities identified the regions associated with SOZ. However, HFO-seq demonstrated a more localized spatial spread over the cortex compared to aHFOs, which is consistent with the occurrence analysis, suggesting a more focused and precise estimation of the EZ (illustrated in Fig. 4A for SOZ and Fig. 4F for resection validation). Among the 25 SZF patients with clinically defined SOZ, the localization error with HFO-seq was significantly lower (2.92 ± 3.56 *mm*, *mean* ± *SD*) than that of aHFOs (18.57 ± 8.9 *mm*), displaying a 818.47% (∼16.58 mm) improvement (*p* = 10^−5^, two-sided Wilcoxon signed-rank test, *n* = 25; Fig. 4B). Additionally, the precision of spatial overlap between the estimated EZ and the SOZ was notably higher for HFO-seq (54 ± 22%, *mean* ± *SD*) compared to aHFOs (45 ± 20%, *n* = 25; *p* = 0.012, two-sided Wilcoxon signed-rank test; Fig. 4C). The recall of overlap with the SOZ was also higher for HFO-seq (77 ± 24%) than aHFOs (71 ± 25%, *mean* ± *SD*; *n* = 25; *p* = 0.016, two-sided Wilcoxon signed-rank test; Fig. 4C). These results suggest that EZ estimations using HFO-seq are more spatially concordant with the SOZ, whereas aHFOs, though relevant, tend to have a broader and scattered spatial activation, which could be attributed to the conflation of various high-frequency activities, possibly including physiological HFOs. Next, by integrating precision and recall into a unified measure, the normalized overlap ratio, which represents the overall concordance between estimation and clinical ground truth, we observed a higher overlap ratio in HFO-seq (63 ± 21%) compared to aHFOs (55 ± 18%, *mean* ± *SD*, *n* = 25; *p* = 0.003, two-sided Wilcoxon signed-rank test; Fig. 4D). Besides, the spatial dispersion also strongly affirmed these findings, showing a smaller spatial spread and greater concordance to the SOZ in HFO-seq than in aHFOs (*n* = 25, *p* = 0.001, two-sided Wilcoxon signed-rank test; Fig. 4E).

Moreover, in a subset of patients who were seizure-free outcome and had post-surgical MRI available, the resection zone was modeled to provide an additional benchmark for clinical validation (Fig. 4F). Specifically, in this cohort, the localization error to the resection zone was substantially lower for HFO-seq (1.31 ± 2.24 *mm*, *mean* ± *SD*) than for aHFOs (16.78 ± 11.33 *mm*), indicating a remarkable deduction of approximately 14.11 mm (*n* = 7, *p* = 0.016, two-sided Wilcoxon signed-rank test; Fig. 4G). The overlap ratio between estimation and resection was slightly better but not significant for HFO-seq compared to aHFOs (*n* = 7, *p* > 0.05), while a notably lower spatial dispersion was observed for HFO-seq (*n* = 7, *p* = 0.016, two-sided Wilcoxon signed-rank test). These results, aligned with previous findings with SOZ validation, suggest that HFO sequences offer a more spatially precise and concordant method for identifying the EZ, contrasting with the broader and more distributed spatial distribution observed with conventional HFOs measure.

### Association of estimated EZ with surgical outcomes

In a subgroup of patients who did not achieve seizure-freedom post-surgery, we sought to explore if our mapping method could differentiate between favorable and unfavorable outcomes. To answer the question, we applied the same mapping technique used in the seizure-free (SZF) group to a cohort of 14 non-seizure-free (non-SZF) patients, and evaluated the results by measuring the localization error, normalized overlap ratio, and spatial dispersion, against the clinically determined SOZ. Upon comparing these metrics between the SZF and non-SZF groups for both HFO sequences and all detected HFOs, we observed a significant deterioration in localization error for HFO-seq in the non-SZF group (7.80 ± 6.79 *mm*, *mean* ± *SD*, *n* = 14), compared to the SZF group (2.92 ± 3.56 *mm*, *n* = 25; *p* = 0.0088, two-sided Mann-Whitney U test; Fig. 4H). Conversely, no marked difference was found for aHFOs (non-SZF: 25.66 ± 10.44 *mm*, SZF: 18.57 ± 8.90 *mm*; *p* > 0.05, two-sided Mann-Whitney U test; Fig. 4H). The normalized overlap ratio showed a remarkable decline of over 34% for both markers for the non-SZF group compared to the SZF group (HFO-seq: *p* = 0.023, aHFOs: *p* = 0.045, two-sided Mann-Whitney U test; Fig. 4I). For spatial dispersion, there was a notable 55% degradation for HFO-seq in the non-SZF group (*p* = 0.02, two-sided Mann-Whitney U test; Fig. S3), whereas for aHFOs, no significant difference was found (*p* = 0.22, two-sided Mann-Whitney U test), though a slight increase (∼11%) was seen in the non-SZF cohort. These results indicate a consistently reduced correlation between the estimated EZ of HFO-seq and the clinical SOZ in patients who did not achieve seizure-freedom.

We further investigated the relationship between the accuracy of HFO-seq localization and surgical outcomes – seizure-free or not. This analysis involved constructing a generalized linear model (GLM) with a binomial distribution, employing a logistic link function (*59*). To account for variations in spatial sampling due to the use of different iEEG electrodes, we normalized the localization measures against the minimum spatial resolution for each patient (see Materials and Methods). In our cohort analysis, we identified significant predictors of surgical outcomes across all considered measures – localization error, overlap ratios, and spatial dispersion, against the SOZ (all *p* < 0.01, Chi-squared test; Fig. S3A). Notably, the precision of overlap emerged as the most reliable predictor (*p* < 10^−5^) with an odds ratio of 0.57 (95% confidence interval (CI): 0.39 −0.85) for non-SZF outcome with each unit increase (Fig. S3A).

Subsequently, the GLMs were employed to forecast seizure outcomes based on the three localization measures, normalized with z-score method for standardization. To ensure balanced representation across different outcome groups, we stratified the training dataset and applied the synthetic minority over-sampling technique (SMOTE) to mitigate class imbalance issues (*60*). The models were trained iteratively on these balanced subsets and evaluated on an independent test set using cross-validation techniques. All models demonstrated substantial discriminative capability, as the receiver operating characteristic (ROC) curves surpassing the chance level performance (Fig. S3B). The model of the highest efficacy, particularly through the overlap precision ratio, yielded an area under the ROC curve (AUC) of 0.9, accuracy of 0.84, precision of 0.95, recall of 0.81, and specificity of 0.91. The evaluation scheme showed a robust and effective performance of the model in predicting seizure outcomes from pre-surgical localization indicators, thereby affirming the predictive validity of HFO sequencing. Overall, these results indicate an inverse relationship between the accuracy of localization measures and the clinical outcomes, in other words, the closer the HFO-seq mapping aligns spatially with the clinically presumed EZ, the higher the likelihood of attaining seizure freedom in patients post-surgery. This correlation might stem from an inadequate investigation of the SOZ during pre-surgical evaluations and highlights the potential utility of HFO sequences in refining EZ localization precision, particularly in scenarios where accurately capturing seizure onset and achieving seizure freedom pose a challenge, offering an additional layer of diagnostic reference to optimize patient treatment strategies.

### Reliability of HFO-seq mapping over long-term recordings

The robustness and reliability of HFO-seq for localization over extended periods were analyzed by examining the variability in localization error across multiple time scales in long-term iEEG recordings. The full recordings were segmented into epochs of varying lengths – 5, 10, 30, 60, and 120 minutes – and the standard deviation of localization error relative to the SOZ was assessed over different recording segments for each patient.

The analysis revealed a significant difference across the different time-scale groups (*n* = 40; *p* < 10^−8^, Friedman’s test; Fig. 4J). Notably, there was a consistent decrease in the standard deviation of localization errors with increasing length of recording segments. For instance, the standard deviation dropped from 4.44 ± 14.95 *mm* for 5-minute segments to 3.94 ± 12.78 *mm* for 10-minute segments, 3.23 ± 10.21 *mm* for 30-minute segments, 2.94 ± 9.73 *mm* for 60- minute segments, and further to 2.54 ± 7.41 *mm* for 120-minute segments (*mean* ± *SD*; all *p* < 10^−4^, two-sided Wilcoxon signed-rank test with FDR correction; Fig. 4J). This trend indicates a stable spatial distribution of HFO-seq measures over time, even with short segments at minute scale. Besides, with a segment of around 30-60 minutes, the mean standard deviation approached the range of the mean localization error. The findings underscore the precision and reliability of the proposed method of HFO-seq mapping in estimating the spatial distribution of epileptogenic activities over multiple lengths of recordings.

### Sequence order and the nodes in relation to the SOZ

Upon the previous observations that the HFO-seq provides marked concordance with the epileptogenic tissue, we sought to explore their spatial origins and relation to the seizure generating areas, or, in other words, whether the sequences typically originate within the SOZ and how they progress at single sequence and electrode level. Specifically, to achieve this, we sorted each HFO sequence by estimating the onset time of each event within the sequence and then categorized the events based on their occurrence within or outside the SOZ (see Materials and Methods), to trace the propagation path of each sequence and interaction with the clinical EZ. A typical example of HFO-seq propagating through multiple electrodes is depicted in Fig. 5A, where the sequence begins at the boundary of the SOZ and traverses through the entire SOZ (additional examples of HFO-seq patterns please refer to Fig. 2 and S2).

**Figure 5.**
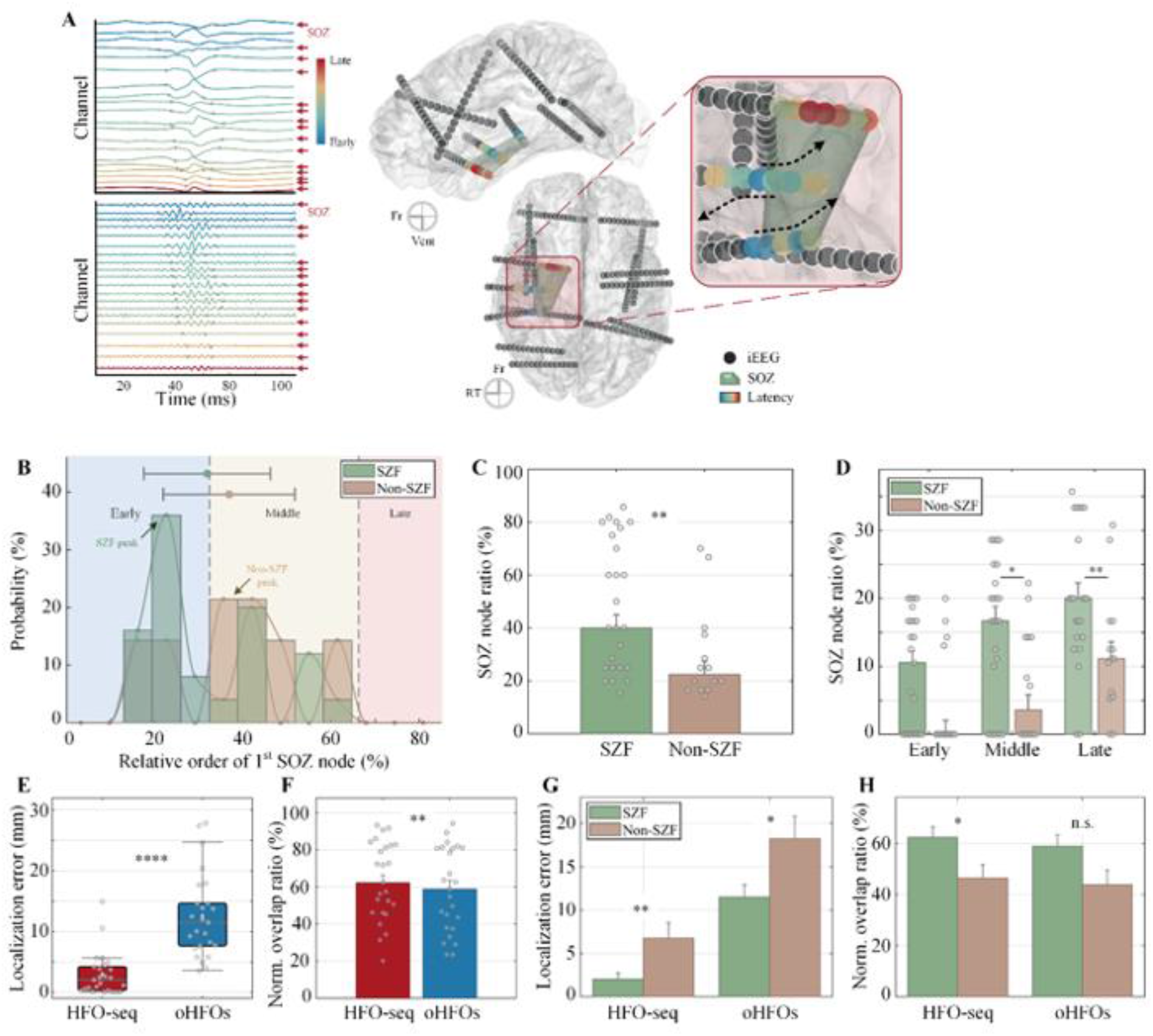
Relative sequence order of HFOs in relation to the SOZ. (A) HFO sequences propagating through multiple channels. A series of HFOs was observed to spread across a localized area, captured by several sEEG electrodes. The latency of each HFO in the sequence is color-coded, from cool (early) to warm (late) colors. The SOZ, located around the hippocampus and amygdala, is marked with red arrows in the left panel and a green volume in the right panel. **(B)** Probability of the relative order of the first SOZ node in a sequence, correlated with seizure outcomes. Three stages are denoted by varying color shades: the first, the middle, and the late one third (∼33%) of HFO sequences. Note the largest peaks of the seizure-free (SZF) and non-seizure-free (Non-SZF) groups. **(C)** Overall ratio between the nodes in the SOZ and the total nodes in each sequence. **(D)** Breakdown of SOZ node ratio into early, middle, and late stages. **(E,F)** The effectiveness of using HFO-seq and onset HFOs (oHFOs) to localize the SOZ is evaluated through (E) localization error and (F) normalized overlap ratio. **(G,H)** The SOZ localization performance, localization error (G) and normalized overlap ratio (H), of HFO-seq and oHFOs in relation to the seizure outcomes. ∗ *p* < 0.05,∗∗ *p* < 0.01,∗∗∗∗ *p* < 0.0001.

First, we examined the relative starting position of the first SOZ node within each sequence for patients with different surgical outcomes. This investigation aimed to understand how the starting points of HFO sequences relate to the SOZ and, by extension, to the underlying epileptogenic network. For patients who had favorable surgical outcomes (seizure-free, SZF), we found that the initial SOZ node marked the highest probability of relative rank at the 23% within sequences, suggesting that in these patients, HFO sequences were more likely to begin near or within the SOZ, reinforcing the notion of SOZ as a critical area in sequence initiation. In contrast, for patients with unfavorable seizure outcomes (non-seizure-free, non-SZF), the initial SOZ node relatively shifted backwards to approximately the 38% mark in the sequences (*p* = 0.33, two-sided Mann-Whitney U test; Fig. 5B), indicating a trend where sequences in non-SZF patients are slightly more likely to start further away from the determined SOZ. These findings suggest that in patients with favorable outcomes, HFO sequences tend to originate closer to or within the SOZ, aligning with previous studies (*43, 44*). Furthermore, we computed the percentage of SOZ nodes in each sequence, revealing a significant disparity between the SZF group (*n* = 25, 49 ± 25%, *mean* ± *SD*) and the non-SZF group (*n* = 14, 30 ± 18%; *p* = 0.01, two-sided Mann-Whitney U test; Fig. 5C). Segmenting a sequence into early, middle, and late stages of equal lengths, we discerned that the primary differences lay in the middle to late stage of the sequences (early: *p* = 0.086, middle: *p* = 0.048, and late: *p* = 0.01, Kruskal-Wallis test with FDR correction; Fig. 6D). This further refined our understanding of HFO-seq dynamics, and suggest that for patients with suboptimal outcomes, the evolution of these sequences and their relation to the SOZ, especially in the later stages, may provide additional insights of epileptogenic tissue localization.

**Figure 6.**
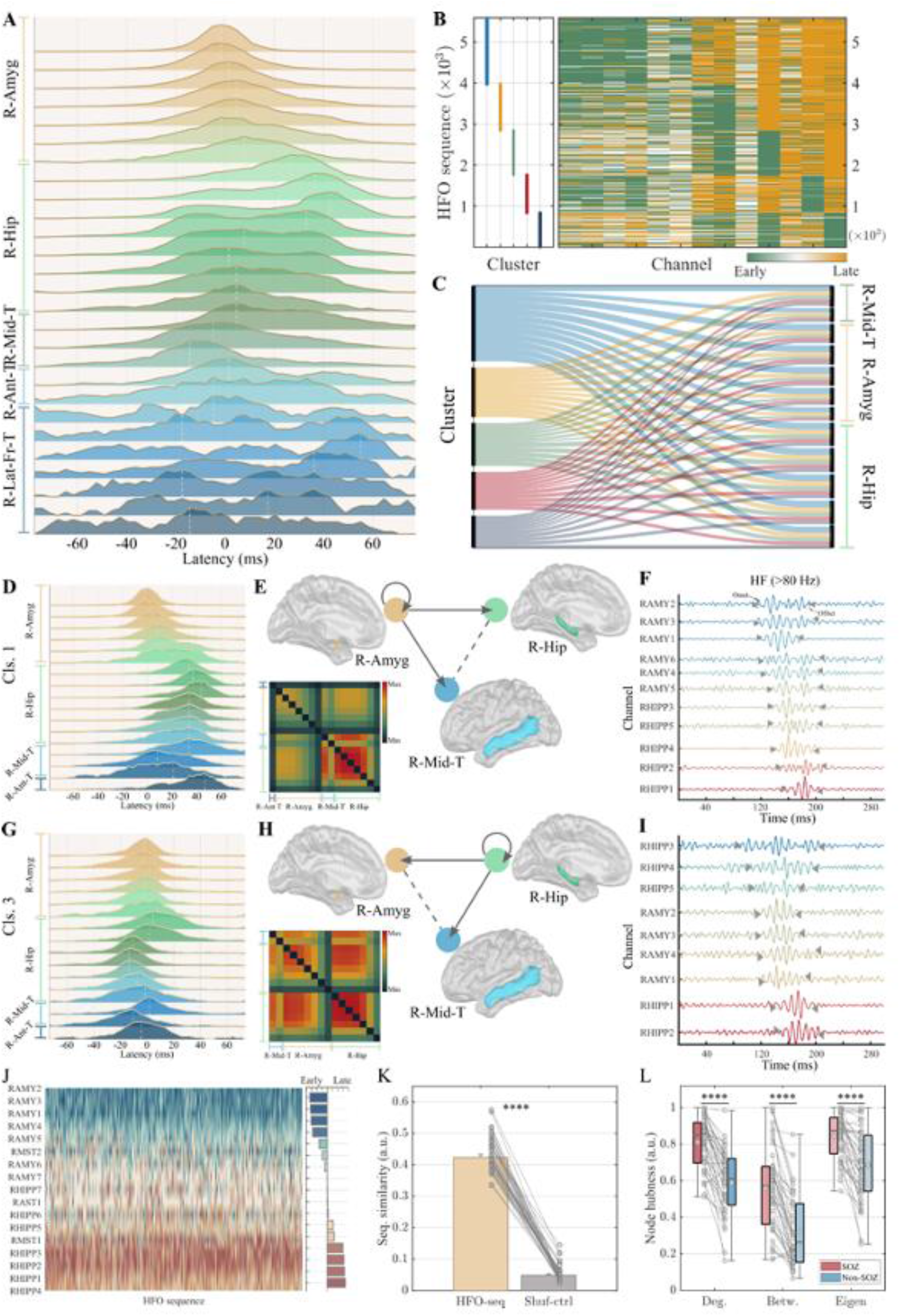
Spatiotemporal dynamics and stability of HFO sequences. (A) Temporal distributions of HFOs onset timings across various channels relative to a specific SOZ channel. Each row represents the probability distribution for one recording channel, with a white dashed line marking the peak probability timing. Notable features include the concentration of channels exhibiting either leading (e.g., R-Mid-T) or lagging (e.g., initial channels in R-Hipp) tendencies, as well as the bimodal distribution consisting of both leading and lagging instances (e.g., later channels of R-Hipp). Also note a flatter, uniform-like histogram from R-Lat-Fr-T. **(B)** Clustering of HFO sequences based on spatiotemporal patterns. The cluster indicies are displayed in the left, with varying colors encoding different groups. The corresponding sequences are enumerated with the sequence indices in the vertical axis and the horizontal axis lists channels engaged in the sequences. The color-coding scheme of HFO sequences represents the propagation tendency of activities within the channels, ranging from cool (early, or upstream) to warm (late, or downstream) colors. **(C)** Association of channels with specific spatiotemporal clusters. Clusters are color coded consistent to the colors in (B) on the left side with the vertical length of each color block proportional the size of the cluster. Channels involved in the sequences are displyed on the right, paired with the respective clusters, and the cortical structures where the channels are located are labeled on the right axis. Note a high degree of spatial overlap in channel involvement across the different clusters. **(D-I)** Two representative and distinct sequence clusters, separating the bimodal channel behaviors noted in (A). **(D,G)** Temporal distriibutions of HFOs onset timings across various channels involved in the sequences of specific clusters. **(E,H)** Sequential interactions among channels within various cortical regions in the identified clusters. The co-activation matrix, color-coded to represent co-occurrence counts, highlights the significant recruitment of the amygdala, hippocampus, and middle temporal regions, aligning with the SOZ. **(F,I)** Examples of HFO sequneces within each cluster, demonstrating different propagation orders, despite considerable overlap in channel recruitment across clusters. The colors of the signals depict the relative timing of onsets, from cool (early) to warm (late). **(J)** Propagation tendency of each sequence within cluster 1 (D-F), with colors encoding the preference for leading or lagging. **(K)** Sequence similarity across all patients against a shuffled control group. **(L)** Hubness measures of nodes in SOZ versus outside (non-SOZ) during HFO sequences: degree centrality, betweenness centrality, and eigenvector centrality. R: right, Amyg: amygdala, Hip: hippocampus, Mid: middle, T: temporal, Ant: anterior, Lat: lateral, Fr, frontal. ∗∗∗∗ *p* < 0.0001.

In addition, we assessed the localization performance using the onset of the propagation HFOs (oHFOs), as established in the literature (*43*), and compared it to entire HFO-seq, examining the concordant with the SOZ in patients with seizure-free outcome (*n* = 25). For oHFOs, the average localization error was 12.51 ± 6.95 *mm* (*n* = 25, *mean* ± *SD*), indicating a reasonably good alignment with the SOZ. However, the analysis of HFO-seq demonstrated a significantly more precise localization, with an error reduced to 2.92 ± 3.56 *mm* (*n* = 25), representing an improvement of 468% (∼9.47 mm; *p* = 10^−5^, two-sided Wilcoxon signed-rank test; Fig. 5E). This finding suggests the superior accuracy of HFO-seq in identifying the epileptogenic tissues. Further analysis on the normalized overlap ratio showed a consistently higher overlap for HFO-seq over the oHFOs measure (*n* = 25, *p* = 0.007, two-sided Wilcoxon signed-rank test; Fig. 5F), while for spatial dispersion, no notable difference was observed (*p* = 0.21, two-sided Wilcoxon signed-rank test).

Moreover, we explored the association of mapping performance with seizure outcomes using oHFOs. This analysis revealed a significant recline in the localization error between patients who were seizure-free (*n* = 25) and those who were not (*n* = 14, *p* = 0.034, two-sided Mann-Whitney U test; Fig. 5G). While the overlap ratio showed a slight increase (∼34%) in the SZF group compared to the non-SZF group, the difference was not remarkable (*p* = 0.15, two-sided Mann-Whitney U test; Fig. 5H). Also, recall that HFO-seq showed generally larger discrepancies in all localization measures between different seizure outcome groups (all *p* < 0.05, Fig. 5G,H).

Overall, the results indicate that the onset of HFOs is a valuable biomarker with good concordance to the SOZ, while analyzing the full extent of HFO sequence, including the propagation patterns, yields more precise localization and higher concordance to the clinical EZ, potentially enhancing the accuracy of surgical planning in epilepsy treatment.

### Multifurcation and spatiotemporal dynamics of HFO sequences

We further delved into the spatiotemporal dynamics and organizational patterns of HFOs as they traverse across cortical areas. This exploration aimed to discern the sequential organization of HFOs and assess the repeatability and stability of their patterns across extended recording periods. For this, the temporal and spatial relationships of individual HFO events within sequences were analyzed and clustered into distinct groups (see Materials and Methods; Fig. 6A-C).

Firstly, we examined the temporal relationships among the channels exhibiting HFO events in the identified sequences, quantifying the onset times of each event and their relative occurrences. An example (Fig.6A) illustrates the co-occurrence timing of channels, aligned to a SOZ channel in the right amygdala (RAMY1). The analysis revealed that while some channels like RMST2-3, consistently led the sequences, others tended to lag (e.g., RHIPP1-2), indicating consistent timing preferences. Intriguingly, certain channels displayed bimodal patterns with instances of leading or lagging (e.g., RHIPP4-6), suggesting variable recruitment orders within the sequences. Besides, uniform distribution patterns in some channels (e.g., right-lateral-frontal-grid) indicated non-time-locked or non-frequent events to the reference channel.

By integrating channel involvement and relative timing as spatiotemporal features, the sequences were clustered into various groups using K-means method (Fig. 6B), identifying a subset of core channels highly involved in HFO propagation. Notably, this HFO-recruited core exhibited considerable spatial overlap with the SOZ, specifically in regions such as the right amygdala, hippocampus, and middle temporal structures (Fig. 6C). Likewise, the channel engagement in each sequence demonstrated similar spatial overlaps and recruitment patterns (Fig. S4). Two distinct sequence clusters were pinpointed, segregating the bimodal channels in the sequences: each with unique initiation and progression paths, though involving similar channels (Fig. 6D-I). Meanwhile, a co-activation matrix, constructed by tallying frequently co-occurring channels within sequences, and community networks, derived using the Louvain method (*61*), also underscored the significant involvement of the amygdala, hippocampus, and middle temporal regions, aligning with the SOZ (Fig. 6E,H). Further analysis of the network properties between the SOZ and external areas, using the hubness of nodes within the co-activation matrix for each patient, revealed higher hubness within the SOZ compared to the outside, indicating enhanced integrally connected nodes to other parts of the network (*n* = 40, all *p* < 10^−6^, two-sided Wilcoxon signed-rank test; Fig. 6L).

Furthermore, we merged the spatial and temporal data to calculate the degree of preference (DP) for each channel within sequences (*46*). This metric represents the tendency of each channel to either lead or lag in the sequences, ranging from -100 (upstream) to 100 (downstream). Note that the HFO sequences illustrate consistent temporal order and unique spatial organization (Fig. 6J). The repeatability of single sequences within each cluster was further assessed by comparing their propagation tendency, using the reproducibility index (RI) to quantify the sequence similarity (*46*). This index is defined as the fraction of consistent channel order compared to a reference sequence determined by the DP across all sequences within a cluster. The analysis revealed a remarkable repeatability of sequences in each patient, significantly surpassing a shuffled control group (HFO-seq: 0.43 ± 0.07, control: 0.05 ± 0.03, *mean* ± *SD*, *n* = 40; *p* < 10^−7^, two-sided Wilcoxon signed-rank test; Fig. 6K). Together, these findings indicate that the organization of HFO sequences involves unique, repeatable, and stable patterns of spatiotemporal propagation engagement of different cortical locations over extended recording periods, offering profound insights into the dynamics of epileptogenic networks.

### Information flow underlying epileptic network constrains the HFOs propagation

We extended our investigation to assess the functional connectivity and directional flow of information that governs the HFOs propagation, asking whether the connectivity differs between the regions involved in HFO sequences and the peripheral areas, examining the intricate spatiotemporal interplay within the epileptic network. This involved analyzing the connections between the SOZ regions and the surrounding areas (non-SOZ), as well as those engaged in HFOs, referred to as the HFO zone (HFOZ), and their counterparts (non-HFOZ), to understand the network dynamics (Fig. 7A).

**Figure 7.**
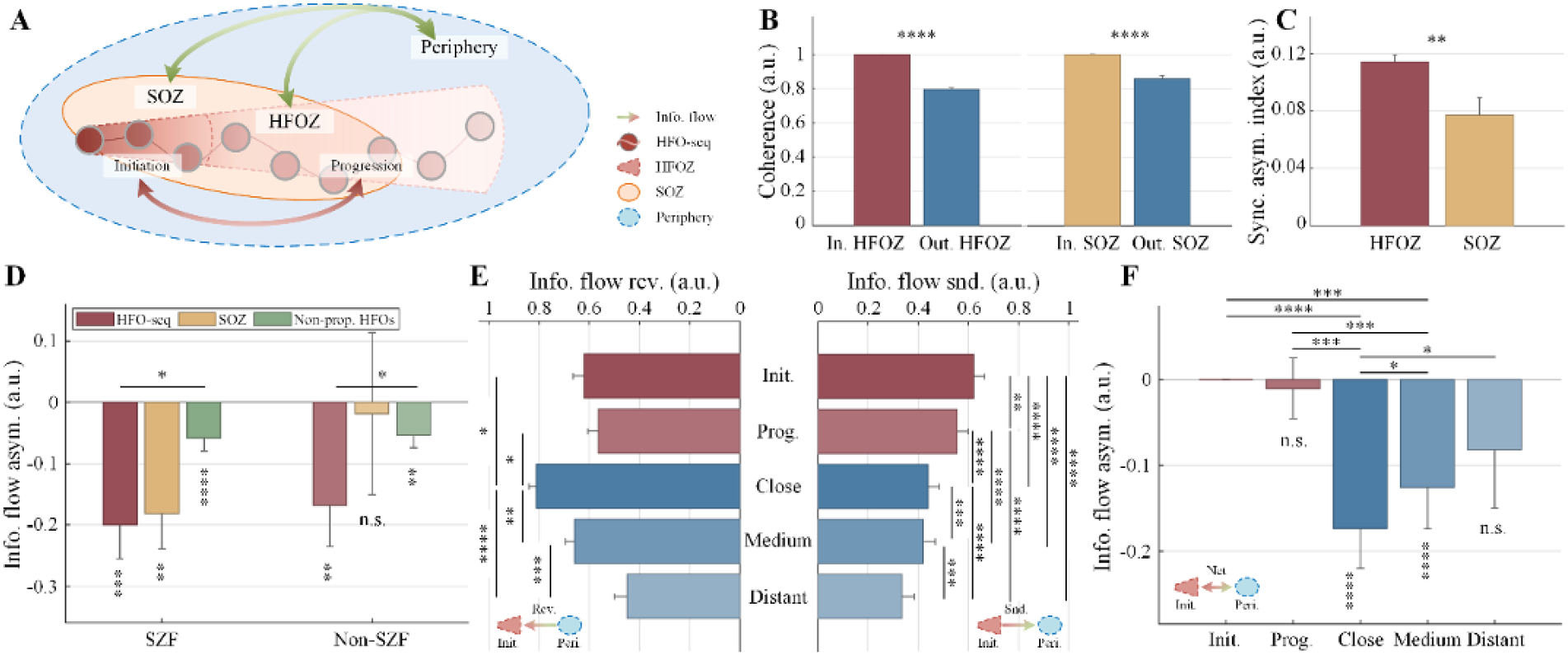
Connectivity of HFOs between foci (SOZ or HFO-zone) and outside. (A) Illustration of connectivity analysis paradigm across SOZ and HFO-recruited zone, as well as the peripheral regions. The HFO-zone is further segmented into the initiation (∼33%) and progression (∼67%) stages. **(B)** Coherence between channel pairs within the HFO-zone (HFOZ) and outside (left panel) and within the SOZ and outside (right panel). Note the higher degree of coordinated activity within the foci compared to outside regions. **(C)** Synchrony asymmetry index between the SOZ and the HFOZ. **(D)** Directional information flow asymmetry between the foci (HFOZ, SOZ) and outside during HFO sequences, compared to a control group of non-propagating HFOs, in relation to different seizure outcomes – seizure- free (SZF) and non-seizure-free (non-SZF). Statistical analyses were conducted to test whether the distributions of each measure, information flow asymmetry, were significantly different from zero, as well as to compare the measured asymmetry between groups. Note a consistent negative information flow asymmetry, indicative of a net inflow of information. This inflow is significantly higher in HFO sequences compared to non-propagating HFOs. However, the information flow asymmetry for the SOZ during HFO sequences were not significantly different from zero in the non- SZF group. **(E)** Information exchange from and to the initiation node with the progression site and the peripheral regions, which is segmented into close (∼15 mm), intermediate (∼55 mm), and distant (∼100 mm) proximity groups. Note the predominant connectivity received from the close proximity to the initiation node and the strong information within the initiation site itself and sending to the progression node. **(F)** Information flow asymmetry between received and outgoing connectivity of the initiation node with the progression site and the peripheral regions. Note significant net inflows from both close and intermediate proximities. ∗ *p* < 0.05,∗∗ *p* < 0.01,∗∗∗ *p* < 0.001,∗∗∗∗ *p* < 0.0001.

Initially, coherence was employed to measure the synchronization of neural oscillations within the foci versus outside. We discovered a notable rise in coherence within the SOZ and the HFOZ, as compared to periphery, indicating elevated neural activity synchronization in both foci during HFO propagation (*n* = 40, both *p* < 10^−7^, two-sided Wilcoxon signed-rank test; Fig.7B). This consistent pattern also highlights a possible interconnection between the HFOZ and the SOZ, also echoed by the observed close spatial overlap between these two regions. Additionally, we calculated the synchrony asymmetry index (SAI) to assess the normalized disparity in coherence between the foci and their surrounding regions, for both SOZ and HFOZ (detailed in Materials and Methods). The analysis revealed a significantly higher SAI in the HFOZ compared to the SOZ (*n* = 40, *p* = 0.008, two-sided Mann-Whitney U test; Fig. 7C), suggesting a nuanced relationship between the networks underlying seizures and HFOs generation.

Moreover, we examined the directional information flow, within and between the SOZ and non-SOZ, as well as the HFOZ and non-HFOZ regions, using the direct transfer function (DTF). Net directional information between the inflow and outflow from the foci was quantified using an information asymmetry index (*62, 63*). For comparison, we analyzed a matched control group of non-propagating HFOs (npHFOs), representing those HFOs being captured by detector yet lacking consistent progression patterns (see Materials and Methods; Fig. S5). Remarkably, a consistent negative information asymmetry (net inflow) towards the foci was observed for HFO-seq and npHFOs (*n* = 40, both *p* < 10^−4^), with a significantly greater inflow for HFO-seq than npHFOs (*p* = 0.0017, two-sided Wilcoxon signed-rank test). Similar net inflow was observed between SOZ and non-SOZ during HFOs propagation (*n* = 40, *p* = 0.0046), suggesting a potential suppression mechanism during propagating HFOs. This trend held true for the seizure-free patient group (*n* = 25; all *p* < 10^−2^ for net inflows and *p* = 0.035 for higher inflow in HFO-seq; Fig. 7D). However, for the non-seizure-free group, this trend only held for HFO-seq and npHFOs (*n* = 14; *p* = 0.02 for HFO-seq and *p* = 0.01 for npHFOs, and *p* = 0.017 for higher HFO-seq inflow; Fig. 7D). Interestingly, the information asymmetry between SOZ and outside did not significantly differ from zero (*p* = 0.43, two-sided Wilcoxon signed-rank test; Fig.7D) in the non-seizure-free group, which might indicate varied networks underlying the SOZ and HFOZ or inconsistency in the spatial definition of the SOZ from the HFOZ in patients with suboptimal seizure outcomes.

Further, to investigate the information flow at a finer spatial scale, we segmented the HFO sequences into initiation (33%) and progression (67%) stages and the peripheral regions into close (∼15 mm), intermediate (∼55 mm), and distant (∼100 mm) proximity groups. We then analyzed the information exchange to and from the initiation nodes within HFO sequences to uncover the potential factors, sequential stage and distance, impacting HFOs propagation. Across all patients, a significant difference in information received by the initiation node from progression region and peripheral regions was noted (*n* = 40; *p* < 10^−5^, Friedman’s test; Fig. 7E). Specifically, the connectivity from the close proximity to the initiation node was the highest, with a significant decrease in information flow among close, intermediate, and distant regions, relative to the distance from the initiation site (*p* < 0.02 for the close node versus the initiation and progression nodes, all *p* < 0.01 for the close node versus the medium and distant nodes, FDR corrected). Meanwhile, the information sending from the initiation site to other nodes differed significantly (*n* = 40; *p* < 10^−20^, Friedman’s test; Fig. 7E), with the highest connectivity observed within the initiation site itself compared to other sites (*p* = 0.0032 for initiation versus progression nodes, and *p* < 10^−6^ for all other peripheral sites, FDR corrected). Similarly, the information from initiation to progression node was remarkably higher than to peripheral regions (all *p* < 10^−4^), and a notable reduction was observed with increasing distance from the initiation site (all *p* < 10^−3^, FDR corrected). We further assessed the information flow asymmetry between received and outgoing connectivity from the initiation node, discovering significant net inflow from both close and intermediate proximities (both *p* < 10^−4^, two-sided Wilcoxon signed-rank test), and the group wise test indicated significant differences (*p* < 10^−4^, Friedman’s test; Fig. 7F). The inflow was substantially higher from the close and medium proximities to the initiation site compared to the progression node (all *p* < 10^−3^, FDR corrected) and the inflow decreased gradually from close to distant regions (all *p* < 0.05, FDR corrected). These findings indicate a strong interconnection within the HFO-zone coupled with suppressions from the surrounding regions that potentially constrain the spread of HFOs, a relationship that is distance-dependent.

## DISCUSSION

This study analyzed the propagation of spontaneous HFOs in patients with focal epilepsy, aiming to advance the delineation of epileptogenic network and to shed lights on the complex spatiotemporal dynamics of HFO sequences and their association with the EZ. Grounded in the hypothesis that the patterns of HFO propagation could serve as precise and reliable markers for identifying pathological networks and delineating the EZ, the research sets out to develop robust methods for mapping the epileptic brain and to explore the propagation mechanisms of HFOs. The study is structured around several key objectives: (i) developing a data-driven approach to identify HFO sequences from iEEG, characterized by a series of HFO events spreading across multiple cortical regions, signify a coherent epileptic network; (ii) establishing the relationship between HFO sequences and the clinically presumed EZs, thereby surpassing conventional HFO measures, and demonstrating the significance of the identified HFO sequences in mapping the epileptogenic tissues and networks, and potentially differentiating seizure outcomes; and (iii) assessing the consistency and reliability of HFO sequences to validate their effectiveness as dependable marker of focal epilepsy; and (iv) investigating the spatiotemporal dynamics of HFO sequences to uncover the underlying mechanisms of HFOs propagation by assessing functional connectivity during these activities.

Together, the primary findings can be summarized as follows. Spontaneous HFOs were observed to traverse across multiple channels in iEEG from epilepsy patients recruited from two clinical centers. These sequences of HFOs, automatically identified from long-term invasive recordings, exhibited a stronger link to the SOZ and lower variability compared to conventional HFOs. Notably, the proposed approach revealed more accurate localization of the clinical EZ through HFO sequences mapping than traditional HFOs, and higher correlation with surgical outcomes. This underscores the potential of HFO sequences as an effective tool for accurately delineating the epileptogenic tissues. Additionally, the reliability of localization was confirmed across extended datasets, with shorter recording segments within one hour providing stable estimations. The investigation also revealed distinctive, repeatable spatiotemporal patterns of HFO sequences, indicating complex dynamics within HFO networks over prolonged observation periods. Furthermore, possible mechanism for the initiation and suppression of HFO propagation was uncovered. It marked a high synchrony of oscillations between the regions involved at the start and throughout the progression of HFO sequences and predominant inward information flows from peripheral to core regions engaged in HFOs, which was observed consistently across different patient outcomes, underscoring its potential importance in the pathology of epilepsy. Interestingly, consistent inward information flows between SOZ and non-SOZ was associated with favorable seizure outcomes, while not for the non-seizure-free patient group. In summary, our research highlights the clinical value and utilities of HFO sequences in epilepsy treatment and management, offering more precise and reliable methods for delineating epileptogenic network and contributing to the broader understanding of epilepsy.

The conventional view and utility of HFOs largely relies on the occurrence rates, with a focus on channels exhibiting on the highest HFO-rate (*28, 31, 64*). However, when defined only by the occurrence, different brain structures may generate activities at variable rates (*65*), and physiological HFOs cannot be disentangled from the pathological ones effectively, which might confound their identification as markers of epileptogenic areas and reduce the specificity of HFOs (*28, 29*). Recent studies have proposed to establish region-specific normative indices for HFOs, thus HFO-rate exceeding a threshold could be considered pathological (*66*). On the other hand, it has also been shown that the HFOs display high variability for the occurrence rates especially when the data segment is short (*31*), which is a critical limitation of prior research, potentially missing the full spectrum of HFO-behavior over time and leading to debatable impact of HFOs in clinical applications (*28, 30, 32*). The present investigation aimed to identify pathological HFOs by incorporating their unique morphological properties and spatiotemporal dynamics. This approach, which focuses on the sequencing of HFO propagation, demonstrates a higher association and better localization to the clinical EZ. This method aligns with studies exploring the propagation of seizure sources and interictal spikes, which have been shown to more accurately localize the epileptic areas (*46, 47, 67*). Our findings suggest that HFO sequences, forming a propagational network, may have greater pathological significance and provide more precise insights into the delineation of epileptogenic tissues and predictive values of seizure outcomes in clinical epilepsy research and practice, offering a more comprehensive and nuanced understanding of epileptogenic networks (*68*).

Previous studies reported that intracranial HFOs propagate in a local extent, indicating that the resection of HFOs onset may be crucial for successful surgical outcomes (*43, 44*). Our findings align with these observations, demonstrating a high likelihood of HFO sequences originating from the SOZ, particularly in patients who achieved seizure-freedom. This suggests a strong correlation between the initiation of HFO sequences in the SOZ and successful surgical intervention. Notably, our research indicates that while oHFOs are indeed valuable biomarkers, a comprehensive analysis of the entire HFO sequences yields a notably more precise and detailed picture for localization of the EZ than merely considering onsets. This is also noteworthy that about 40% of sequences may initiate outside of SOZ, even in seizure-free patients. For those with suboptimal outcomes, the clinically defined SOZ nodes tend to be recruited in the later phase of HFO sequence, which further underscores the complexity of epileptic networks and the variability in how they manifest in different patient groups. A notable limitation of earlier studies is their reliance on short data segments, potentially missing certain variations in HFO propagation patterns. This also suggests the participation of regions not identified or examined as part of the SOZ in the genesis of these sequences, which might be involved in the broader pathological network, especially in challenging cases with unfavorable outcomes. Meanwhile, significant differences in the SOZ involvement in sequences were observed, especially in the later portions of sequence nodes, suggesting a dynamic interaction between HFO propagation and seizure network, which may imply evolution of epileptic network over time, perhaps reflecting changes in neural connectivity or plasticity (*69*). It also suggests that the epileptogenic network may extend beyond the traditionally defined SOZ or suggesting the SOZ, as currently delineated, might not fully capture the extent of the seizure network, especially for patients with unfavorable outcomes in this study. The findings may provide additional reference for refining the extent of epileptic networks in these challenging cases.

This study expands upon previous research by analyzing extended recordings, diverging from the typical focus on brief data segments (*43, 44*). Such analysis is pivotal, as prolonged recordings are suggested to be essential for sampling variabilities and deriving robust conclusions about HFOs (*31*). However, collecting and analyzing long-term data in clinical settings can be challenging, especially in the absence of automated tools, considering the cost of expert review (*33, 34*). Prior studies often relied on short segments of intracranial recordings for HFO detection, which could lead to significant variability and even conflicting results in conclusions about HFOs and their spatial localization (*30–32*). Our study addresses this limitation by employing an extensive HFO detection methodology in long-term recordings, analyzing millions of HFOs. This approach not only confirms earlier findings that conventional HFO rates vary in spatial localization (*31*), but also demonstrates the stability of HFO sequences over different data lengths. Our findings indicate that, despite possible variations, regions frequently recruited by HFOs propagation may possess significant epileptogenic value, suggesting that HFO sequences are stable characteristics of the epileptic network rather than transient events, bearing implications as a refined interictal biomarker for surgical planning and epilepsy treatment, potentially reducing the duration of invasive recordings and improving epilepsy management.

Moreover, we delved into the intricate propagation patterns of HFOs, revealing diverse spatiotemporal dynamics across individual patients. This aspect of HFO research, where initial studies focused primarily on the initial onset and the congruence with SOZ (*43, 70*). is relatively untapped. Consistent with earlier findings, our analysis indicates that SOZ nodes often trigger the initial phase of HFO sequences. However, our results extend beyond this, revealing that HFO sequences can also emerge from the periphery of SOZ or even beyond it, forming multifurcation morphologies, while the highly-involved core of HFO sequences exhibited high coincidence with the SOZ, especially in patients with favorable seizure outcomes, which might suggest the critical role of overlapped regions in traversing pathological activities forming epileptic network. The intricate interactions within the HFO network likely play a crucial role in the observed variability in the initiation of HFO sequences. In addition, our study underscores that the cortical organization of HFO sequences is characterized by distinct yet repetitive and consistent patterns of either spatial or temporal propagation, engaging different cortical locations across extended periods. Recent research indicates that epileptic tissue may be transiently impaired by pathological activities (*39, 71, 72*), as opposed to being entirely dysfunctional. This idea aligns with the observations of HFOs activations being linked to a range of hypersynchronous states and inhibitory processes (*14, 73, 74*), such as the reduced inhibition in pathologically interconnected neuron clusters, generating pathological HFOs (*75*). We speculate that such transient impairment of epileptic tissue could lead to diverse network disruptions, accounting for the variety of spatiotemporal HFO patterns within the identical epileptic network.

Our investigation into the functional connectivity of cortical regions involved in HFOs propagation uncovered significant neural synchronization and patterns of information flow. We observed substantial coherence, indicating enhanced neural oscillation synchrony within both the SOZ and the HFOZ, with coherence being higher in the HFOZ, which suggest inherent relationship between the cellular and network mechanisms of seizures and HFOs(*26*) and might be consistent with hyperexcitability linked to fast-spiking interneurons (*76*), which may play a crucial role in the initiation and spread of seizures. These results suggests a nuanced relationship between these regions, highlighting both shared characteristics and distinct differences in HFO generation (*26, 52, 77*).

We also observed a pronounced inward flow of information in areas associated with HFO sequences, suggesting a suppressive effect from surrounding neurons that may inhibit the spread of HFOs. This suppression could be linked to inhibitory synaptic mechanisms, such as the activity of GABAergic interneurons, to control excessive excitability (*73, 76*). Considering the substantial spatial overlap between the HFOZ and SOZ in patients with seizure freedom, the consistent inward flow in both zones points to similar underlying networks (*26*). The variance in the intensity of inward flow between propagating and non-propagating HFOs indicates different levels of inhibition required to mitigate their influence on adjacent tissues, echoing research on seizure propagation and cessation (*62*), which could be variable depending on the strength of pathological activities and level of recruitment of the local extent.

Moreover, the distinct information flow patterns between initiation and progression areas compared to peripheral regions reveal a tight interconnection within the HFO-zone, moderated by external suppression. This dynamic, coupled with observed elevated synchrony during HFO propagation, suggests a complex mechanism of excitation-inhibition balance regulating HFO initiation and progression, aligning with previous studies on push-pull antagonism and its critical role in epileptic networks (*62, 63, 78–80*). Alterations in excitation-inhibition balance could significantly influence the development of interictal activities and evolvement into seizures (*62, 78*), highlighting the critical role of disruption and/or recalibration in synaptic transmission dynamics involving both excitatory and inhibitory neurotransmitters, like glutamate and GABA, in brain function and pathology.

Interestingly, differences in information flow patterns within the SOZ of patients who did not achieve seizure freedom post-treatment, as compared to those who did, might provide additional evidence of inconsistency in the clinical SOZ and reinforce predictive value of HFO sequences. The discrepancies echo the findings in localization and morphology analysis, pointing to the possible involvement of regions not traditionally identified or examined with the SOZ but significantly recruited by HFO sequences, receiving major suppressing effect from the periphery, may of importance. These results consensus on the potential value of predicting seizure outcomes using connectivity methods (*81*), and underscore distinct characteristics of HFO sequences in patients with suboptimal outcomes, suggesting that regions beyond the conventional SOZ play a critical role (*63, 82*).

In this research, we present a systematic approach to analyzing HFO sequences in iEEG recordings for delineating the epileptogenic network in epilepsy patients. Our study advances the understanding of complex neural activities associated with epilepsy by developing an HFO- sequencing technique that automates the identification and sequencing of pathological HFOs, offering enhanced precision and deeper insights compared to conventional HFO analysis, promising to improve epilepsy surgery planning and outcomes for complex cases. Clinically, the HFO-seq method emerges as a crucial tool for mapping the EZ in long-term iEEG recordings, serving as a robust and refined interictal biomarker. Its application could potentially shorten the duration of invasive procedures, facilitate non-seizure monitoring, aid in therapy evaluation, guide surgical planning and implantation, and assist in assessing disease severity, therapeutic efficacy, and post-operative evaluations. Diverse evidence from localization, spatiotemporal dynamics, and connectivity analyses reveals distinct patterns in patient cohorts with varying seizure outcomes, underscoring the predictive value of HFO-seq and provide additional insights for better delineating the epileptogenic networks in patients with challenging conditions. The multifurcation and connectivity analysis uncover a complex interplay of neural activities, indicating a critical excitation-inhibition balance in regulating HFOs spread. These insights open doors to more targeted and effective therapeutic approaches for epilepsy management, urging further investigation into the dynamics and mechanisms of epileptogenic networks. A thorough and longitudinal understanding of HFO dynamics could significantly refine the identification of epileptogenic zones and surgical success rates. Ultimately, incorporating these findings into clinical practice could lead to more sophisticated diagnostic tools and refined surgical strategies, especially for patients with complex epileptogenic networks.

While our study primarily focuses on focal epilepsy, the methodologies developed hold potential applicability to various epilepsy types, including those with ambiguous seizure origins. The limited spatial coverage of iEEG electrodes and partial covering of epileptogenic tissues represent significant challenges in epilepsy monitoring. Source mapping with invasive recording could enhance localization accuracy (*67, 83*), yet it is influenced by the listening field of recording contacts (*84*). Combining intracranial and scalp recordings may offer improved source localization accuracy, especially for deep brain structures, necessitating advanced signal processing and data integration techniques (*85*). In addition, the spatial resolution of current investigation using macro- electrodes is limited at millimeter scale, finer recordings with micro-electrodes might be necessary to further indicate the detailed mapping and investigation of HFOs network (*86*). Moreover, investigating the cellular and neurophysiological mechanisms behind HFOs generation, propagation, and interaction with variable spatiotemporal dynamics within the epileptic network remains a promising research avenue. Expanding HFOs research across various epileptic conditions will realize their utility in epilepsy management. Translating knowledge on HFOs and network properties into clinical practice could significantly improve the management of epilepsy, offering new diagnostic and monitoring tools and the enhancement of surgical outcomes for patients with epilepsy, benefiting a broader range of epilepsy patients.

## MATERIALS AND METHODS

### Study design

The primary objective of this study was to develop an integrated approach for identifying spontaneous HFO sequences and precisely delineating epileptogenic tissues and to investigate their spatiotemporal dynamics and stability in patients with drug-resistant focal epilepsy. We collected a dataset from 40 retrospective patients who underwent long-term invasive EEG monitoring and met inclusion criteria at Mayo Clinic (Mayo, Rochester, Minnesota) and University of Pittsburgh Medical Center (UPMC, Pittsburgh, Pennsylvania). Comprehensive presurgical assessments were conducted, including medical history, neurological and neuropsychological examinations, high-resolution pre-implantation MRI, post-implantation CT, and long-term iEEG recordings. No data was excluded from this analysis. The study was approved by the Institutional Review Boards (IRB) at Carnegie Mellon University, Mayo Clinic, and the University of Pittsburgh. All patients gave informed consent to participate in this study. The standard clinical care of patients was not influenced by their participation. Clinicians involved in data collection and marking clinical findings were blinded to group allocation and data analysis results. Investigators were not blinded to clinical information. Patient-specific brain models were constructed based on high-resolution MRI scans of individual patients. Millions of HFOs identified from the prolonged recordings were sequenced and analyzed using our proposed automatic and semi-automatic techniques. The identified HFO sequences were evaluated using established measures, including occurrence rates and localization metrics, in comparison to clinically defined evidence, such as SOZ and resection volume. This assessment was benchmarked against published HFO-related markers. The spatiotemporal propagation of sequences across cortical regions was analyzed to assess the stability of HFO sequences and to gain insights into their distribution patterns, repeatability, and functional connectivity. Patients were grouped based on the availability of clinical information as well as the outcomes of surgical intervention (i.e., seizure-free and non-seizure-free), according to the international league against epilepsy (ILAE, Mayo) and Engel classification (UPMC) (*87, 88*) by clinicians, providing standardized measures of post-surgical success in epilepsy treatment.

### Patient population and data collection

The study included 40 patients (24 females, aged 14-60 years) with drug-resistant focal epilepsy. All patients were recruited and evaluated by expert epileptologists at Mayo Clinic and UPMC. The inclusion criteria were detailed as follows: (i) Undergoing complete pre-surgical investigations, including high-resolution MRI and CT, long-term iEEG recording (over 24 hours), and neuropsychological assessment; (ii) Presence of a non-generalized SOZ and eligible for surgical intervention; (iii) Availability of iEEG recordings with minimal artifacts and a clearly identified SOZ, as determined by clinicians; and (iv) A minimum one-year follow-up period post-surgery, enabling the evaluation of seizure outcomes.

All participants underwent extensive intracranial pre-surgical monitoring using a digital, 256-channel Xltek system (Natus Medical Inc., California, USA), with the reference electrode generally positioned on the dura mater away from the epileptic field. The data were recorded with a sampling rate of 500 Hz for Mayo Clinic dataset and 1-2 kHz for UPMC dataset and high pass filtered above 1 Hz to remove spurious slow activity and DC drifts. To align with the Mayo Clinic dataset, the UPMC data was down-sampled to 1 kHz, using an anti-aliasing filter set at 250 Hz. Structured MRI scans were conducted in line with a standardized seizure protocol (Prisma 3T or mMR, Siemens, Munich, Germany; SIGNA 3T or GE Architect or GE Discovery MR750, GE, Boston, MA, USA). These scans were performed for pre-surgical planning and for post-operative evaluation, when available. Co-registrations of post-implantation CT and post-resection MRI with pre-surgical MRI were conducted. This process was essential for creating three-dimensional models of both the iEEG electrodes placement and the resection zone. Accurately defining these models was crucial for delineating the clinical EZ and for the comprehensive evaluation of mapping results. The post-surgical outcomes were determined by the clinicians following the ILAE and Engel classification (*87, 88*), during the follow-up period (over 12 months), and patients with ILAE 1-2 and Engel I outcomes classified as seizure-free, while others as non-seizure free outcome.

### Statistical analysis

Statistical details are provided in the Results and figure legends. Descriptive statistics are shown as *mean* ± *SD*, unless specifically indicated. Statistical significance was set for ∗ *p* < 0.05. To address multiple comparisons, the false discovery rate (FDR) method was employed, ensuring a 5% control threshold for false discoveries. All statistical analysis and visualization were conducted in Matlab.

## List of Supplementary Materials

Materials and Methods Fig. S1 to S5 References

## Acknowledgments

The authors gratefully acknowledge all patients involved in the study and thank Drs. Haiteng Jiang, Abbas Sohrabpour, and Rui Sun for useful discussions on data analysis and help in data preparation. We thank Drs. Vasileios Kokkinos, Boney Joseph, Andrea Duque, and Shuai Ye, as well as Fan Yang for assistance in data preparation, and Cindy Nelson in data collection. This work used the Extreme Science and Engineering Discovery Environment (XSEDE), which is supported by NSF grant ACI-1548562. Specifically, it used the Bridges-2 system, which is supported by NSF award number ACI- 1928147, at the Pittsburgh Supercomputing Center (PSC).

## Author contributions

Conceptualization and methodology: ZC, BH; Data curation: MR, GAW, AB, XJ; Formal analysis and visualization: ZC; Investigation: ZC, BH; Supervision: BH; Writing – original draft: ZC; Writing – review & editing: ZC, BH, MR, GAW.

## Data and materials availability

All data supporting the findings of this study are available within the article and its supplementary materials. The analysis codes with de-identified sample data will be made public upon paper acceptance. The developed scripts and codes were written and tested using Matlab (version 2018a, 2023a). We employed Curry (version 7-9, Compumedics, NC, USA) and EEGLab toolbox (version 13.6.5b, https://sccn.ucsd.edu/ eeglab/index.php) for pre-processing analyses and visualization. Additionally, open source toolboxes were used for signal processing and statistical analysis purposes, including the FieldTrip (https://www.fieldtriptoolbox.org/) and FDR toolbox (https://www.mathworks.com/matlabcentral/fileexchange/27418-fdr_bh).

