## Supplementary_material for "Spontaneous HFO Sequences Reveal Propagation Pathways for Precise Delineation of Epileptogenic Networks"

**List of Supplementary Materials**

Materials and Methods

Fig. S1 to S5

References

### Supplementary Materials:

### Materials and Methods

#### Detection of interictal high-frequency oscillations

Long-term intracranial electroencephalography (iEEG) recordings were obtained for each patient, extending up to 27 hours, during interictal periods. These recordings were captured at a minimum sampling rate of 500 Hz. The length of data used for analysis varied, depending on the availability and quality of the continuous clinical monitoring, with an ideal target of approximately 24 hours per patient. Initially, a visual examination of the recordings was conducted to verify data quality and to ensure minimal contamination by artifacts. Data from ictal periods, as identified by clinicians, were excluded from analysis. Any segments with evident artifacts or saturation were also removed. Subsequently, the raw iEEG data were high-pass filtered above 1 Hz to remove baseline drifts and notch filtered at 60 Hz, including harmonics, using an FFT Hann filter with a slope of 2 Hz. The sampling rate for Mayo Clinic dataset was 500 Hz; for UPMC dataset, the sampling rate varied between 1 to 2 kHz; thus, to ensure consistency, the data was standardized and down sampled to 1 kHz, with an anti-aliasing filter set at 250 Hz. The recordings were then re-referenced; for electrocorticography (ECoG) data, a common average reference was employed *(1, 2)*, and for stereo-electroencephalography (sEEG) data, a low-variance white matter reference was utilized *(3)*. Automated screening of extracted long-term iEEG data was conducted using a previously established method for high-frequency oscillations (HFOs) detection *(4)*, and each channel was processed independently. For each patient, the raw EEG data underwent initial filtering with a zero-phase FIR band-pass filter ($64^{th}$ order) from 80 to 240 Hz. This step was followed by an energy detector screening to identify potential high-frequency activities (HFAs) distinct from background noise. Specifically, in each channel, a 100 ms moving window was used to calculate the standard deviation distribution of the signal amplitude. A baseline threshold was then set at five times the median of the distribution. Any sample in the filtered signal exceeding this amplitude threshold was marked as an initial candidate for HFOs. Generally, a low baseline was employed to ensure high sensitivity in this phase.

Each detected candidate was analyzed over a 256 ms segment, comprising 128 ms before and after its peak. Within this segment, the background envelope of the first and last 80 ms of the filtered epoch was calculated using the Hilbert transform, and a baseline was set at three times the median of the envelope distribution. Then, the number of crossing relative to this baseline was counted for the central 80 $ms$ for each detected event and those events with crossings exceeding eight (which equates to two per each of the four cycles) were selected from the initial detections, aligning with the clinical definition of typical HFOs *(5, 6)*. Besides, the number of zero-crossings in the unfiltered signal was also assessed, and those with more than ten crossings were discarded, as they were likely attributable to noisy activities *(7)*.

To discriminate the putative HFOs with high precision, we employed a comprehensive set of features, incorporating aspects of temporal, spectral, and spectrogram analysis. These features were extracted from each event that survived initial screening, in both unfiltered and filtered data (above 80 Hz). Time-frequency representation of these detected high-frequency events in the raw data was utilized to define the characteristics of each event across both low and high-frequency domains. This method included three categories of patterns: temporal, spectral, and spectrogram, as previously established *(4)*. Subsequently, the extracted features facilitated the segregation of detected events into distinct clusters using unsupervised Gaussian Mixture Model, initialized with k-means clustering. The optimal number of clusters was identified using the elbow method *(8)*. Upon clustering, the characteristics within each cluster were visualized through piled waveforms of both unfiltered and filtered data, along with the spatial distribution of channels detecting these events (Fig. S1). The morphology of traces, both unfiltered and filtered, aided in differentiating clusters of putative HFOs from artifacts. As illustrated, various HFOs clusters were observed with distinct repetitive patterns, aligning with the previous findings *(4, 9)*. The consistency of these clustered events was verified by examining the repetitiveness in the stacked signal waveforms and the spatial channel distribution. Those clusters containing evident artifacts, such as strong transient or high amplitude artifacts (Fig. S1), were excluded. The remaining events were preserved as putative HFOs for further analysis.

**Identification of HFO sequences**

All detected HFOs from the initial phase were aggregated and categorized based on their temporal and spatial characteristics, namely, their occurrence time and channel location. We defined a "HFO sequence (HFO-seq)" as a series of HFO events spanning multiple electrodes within a brief timeframe, representing a spatiotemporal co-activation or propagation of HFO activities. These sequences were identified by analyzing the spatiotemporal patterns of the aggregated HFOs during long-term monitoring using our proposed approach, which is constructed based on a previously established procedure for interictal spike analysis with modifications *(10–12)*.

The identification process commenced by selecting an initial HFO event as the "leading event" of a potential sequence. Subsequent HFO events occurring within a 150 ms window of this leader were incorporated into the sequence. Besides, any events within 15 ms of an existing sequence member were also appended. This process was iteratively repeated, selecting the next available event as the leading event, until all HFOs were evaluated, and sequences identified. Subsequently, we applied a set of criteria to filter out sequences of lesser significance. Firstly, sequences comprising fewer than three events were automatically discarded. Considering the plausible speed of neural transmission, we imposed a maximum propagation velocity limit of 10 m/s between any two adjacent HFO events within a sequence *(13, 14)*. Exceptions to this limit were permissible if frequent co-occurrences (exceeding 5% of the quantile of connections at the electrode level) suggested rapid propagation scenarios, such as through gap junctions. Sequences where over half of the events were temporally clustered within a 2 ms window were excluded, as they likely resulted from common artifacts across channels. Additionally, a clustering method was employed to further refine the sequences, specifically removing the occasional sequences that demonstrated less similarity to others, based on the features derived from trajectory mining techniques *(15)*. The sequences that successfully met these criteria were classified as putative HFO sequences. This sequence identification step plays a key role as a discerning tool for pathological HFOs. It is based on the observation that when HFOs occur concurrently within a network in a coordinated spatiotemporal manner, they are more likely to be closely associated with underlying pathological processes.

**HFO rates and asymmetry measure**

A common method for quantifying the activation of HFOs involves calculating their occurrence rate at either the sensor or regional level. This method provides a measurable indicator of HFO activity, facilitating comparisons and analyses across different spatial domains of brain activity. Interpretations of HFOs typically rely on channels exhibiting the highest rates of HFO activity *(16–18)*. The "occurrence rate" is defined as the number of HFO occurrences per minute within a given channel. Another metric of interest is the "HFO rate asymmetry," which represents the normalized difference between average HFO rates inside and outside the seizure onset zone (SOZ). This metric provides insights into the correlation between HFO rates and the SOZ as established by clinical findings *(16, 19)*.

$$R_{asym} = \frac{r_{SOZ} - r_{NSOZ}}{r_{SOZ} + r_{NSOZ}}$$

where $r_{SOZ}$ is the average HFO rate inside the SOZ and $r_{NSOZ}$ is the rate outside. The asymmetry measure, a normalized value ranging between $-1$ and 1, facilitates an intuitive and retrospective interpretation of the relationship between the HFO rate and the SOZ or other clinical evidence. Specifically, a negative asymmetry value suggests a predominance of HFOs outside the SOZ, whereas a positive value indicates a higher concentration of HFOs within the SOZ, potentially signifying clinical relevance. This asymmetry index is commonly used as a normalized metric, which quantifies the degree of asymmetry or imbalance between two comparable entities, and was further applied in subsequent spatiotemporal and connectivity analyses.

**Mapping of HFO sequences and performance assessment**

For each patient, an individualized cortical model was constructed using segmentation of gray matter from their presurgical magnetic resonance imaging (MRI), using Brainstorm *(20)* and Curry 9 (Compumedics, NC, USA). The modeling of iEEG electrodes, encompassing both grids and depth electrodes, was based on their three-dimensional locations. These locations were co-registered to the pre-surgical MRI and then projected onto the individualized cortical model. Similarly, the area of surgical resection was segmented from the post-surgical MRI, and then projected onto the cortical model as part of the co-registration process between the post-surgical and pre-surgical MRIs for individual patients.

To estimate the overall cortical distribution of HFO sequences over a specified timeframe, the sequences were cumulatively mapped as trajectories across their spatial extents. The activation patterns were analyzed by evaluating the spatial distribution of HFO-seq across all sampled locations of the iEEG electrodes. The extent of the active regions of HFO sequences, also referred to as the highly recruited HFO-core, was determined using a threshold derived from the 10% confidence interval of the mean spatial distribution, calculated through a bootstrap method with 1000 replications *(21)*. This technique provided an overarching estimation of the EZ using HFO-seq data. Similarly, this ampping approach was also applied to analyze other well-established benchmarks, including all detected HFOs (aHFOs) and the onset of HFOs (oHFOs), to estimate their respective spatial distributions. The estimated distributions of these HFO markers were quantitatively evaluated for each patient by comparing them against the clinically presumed epileptogenic zone (EZ), which was identified based on key clinical findings, including the SOZ and the areas of surgical resection. The SOZ was localized through the co-registration of post-implantation CT images with pre-surgical MRI, as determined by clinical experts, and the resection area was modeled from post-surgical MRI, co-registered with pre-surgical MRI. To conduct this evaluation effectively, several established performance metrics were utilized: (i) localization error (LE), (ii) normalized overlap ratio (NOR), and (iii) spatial dispersion (SD) *(22–25)*. The localization error (LE) was defined as the minimum distance between the identified HFO markers and the nearest point in the clinically defined EZ (the "ground truth"). The normalized overlap ratio (NOR) was calculated by determining the overlap between the estimated distribution and the clinical EZ, normalized by the area of either the ground truth or the estimation. This ratio was expressed in two forms, recall and precision, and the combined geometric mean was also computed to provide an integrated value ranging from 0 (no overlap) to 1 (perfect overlap). Spatial dispersion (SD) was quantified as the weighted quadratic mean of the minimum pairwise distances between the estimated distribution and the clinical EZ. This metric characterizes the spatial spread of the estimated distribution relative to the ground truth. The specific mathematical formulations for calculating the normalized overlap ratio and spatial dispersion are detailed below.

$$NOR = \sqrt{\frac{S_{Ovlp}^2}{S_s S_G}}$$

where $S_{Ovlp}$ is the overlap area between the estimation and the ground truth, and $S_s$ and $S_G$ are the area of the estimation and the ground truth, respectively.

$$SD = \sqrt{\frac{\sum_{i=1}^{N_s} d_{i,G}^2 \hat{j}_i^2}{\sum_{i=1}^{N_s} \hat{j}_i^2}}$$

where $\hat{j}_i$ is the spatial recurrence rate of estimation $i$, $d_{i,G}$ is the minimum distance from the estimation $i$ to the ground truth, and $N_s$ is the number of distributed locations in the estimation.

**Spatiotemporal dynamics and repeatability analysis of HFO sequences**

To comprehensively analyze the spatiotemporal dynamics of HFO sequences, we focused on quantifying the organization of channels involved in the sequences and the temporal order in which events appeared in each channel. Spatially, we mapped the dynamics by pinpointing the three-dimensional locations of channels implicated in each sequence. Temporally, we measured the onset and offset timing of each event, calculating the first and last samples that crossed the envelope threshold, which was defined by the 95% confidence interval of the mean amplitude distribution, determined using a bootstrap technique *(21)*. The events within each sequence were then chronologically ordered based on their onset times. We developed a method to encapsulate both spatial and temporal characteristics of these sequences within a unified matrix, to represent the sequential order of channel involvement in each HFO sequence. Specifically, we categorized each HFO sequence into the upstream, the intermediate, and the downstream part. The involvement order of channels in each segment was mapped onto an array using -1 for upstream events (indicating earlier involvement), 1 for downstream events (indicating later involvement), and 0 for intermediate events or channels not involved in any event. By concatenating these arrays, we constructed a matrix that effectively signifies the temporal flow of HFO activity within each channel, thus distinguishing between upstream and downstream tendencies. This matrix provides a detailed view and representation of HFO network dynamics and lays the groundwork for subsequent investigations into the spatiotemporal arrangements of HFO sequences and the stability of such properties. Utilizing the spatiotemporal property matrix, we applied the k-means clustering method to categorize the HFO sequences into multiple clusters, with the optimal number of clusters determined using the elbow method *(8)*. This clustering step aimed to distinguish various HFO propagation patterns, which were characterized by their distinct spatial and temporal preferences, effectively leading to a multifurcation of HFO sequences, as depicted in Fig. 6A-C. Effectively, distinct clusters with unique patterns illustrate differentiated bimodal channels within these sequences. Also note that though each cluster demonstrated specific and varied initiation and propagation paths, these clusters involved similar channels overlapping spatially, recruiting similar cortical regions during the propagation of HFOs as illustrated in Fig. 6D-I.

Besides, a co-activation matrix was constructed by counting the occurrence of channel pairs listed across all HFO sequences. Specifically, for each HFO sequence, the channels involved were considered co-activated, and each pair of these channels contributed to incrementing the corresponding element (channel pair) in the matrix. Consequently, the co-activation matrix is symmetric (Fig. 6E,H). Notably, this matrix highlighted the significant involvement of key brain regions. In the representative example, the presence of HFO activity in regions such as the amygdala, hippocampus, and middle temporal regions is consistent with the SOZ, underscoring the pathological significance of these specific brain regions in the context of HFO activity.

To further elucidate the regional propagation dynamics of each channel in HFO sequences, we computed a "degree of preference" (DP) measure *(12)*. This process began by tallying the number of upstream and downstream events for each channel across all HFO sequences, and DP was then defined as the percentage ratio, calculated from the difference and sum of downstream and upstream events in each sequence. This metric effectively demonstrates the spatial propensity of a channel to either precede or follow in the temporal propagation of an HFO sequence, with its values ranging from -100 (indicating consistent upstream activity) to 100 (signifying consistent downstream activity). In an illustrative example (see Fig. 6J), the sequential order of each HFO sequence is depicted along with the upstream/downstream tendency of each channel, revealing highly repeatable temporal propagation patterns and a unique spatial organization of HFO events.

Moreover, we assessed the repeatability of sequences within each cluster by comparing the propagation tendencies of each channel across all sequences. The reproducibility index (RI) was employed for this purpose *(12)*, defined as the proportion of consistent elements in each sequence compared to a reference sequence. The reference sequence was determined based on the DP to outline the overall propagation order across all channels. RI values range from -1 (indicating a complete mismatch) to 1 (denoting a perfect match). For each patient, RI was calculated for each sequence cluster and subsequently weighted by the size of the clusters. To establish a control for comparison, we created a randomized group by performing random permutations of the channel orders within each sequence while maintaining the original timing of events. This process resulted in a shuffled sequence dataset, serving as a benchmark to validate our findings. This approach allows for an assessment of the overall similarity and repeatability of the sequence patterns. Collectively, these employed methods were designed to investigate the cortical organization of HFO sequences.

**Connectivity analysis of HFO sequences propagation**

To investigate the connectivity among regions implicated in HFO sequences progression, we implemented various analytical methods to assess network connectivity and explore the driving mechanisms behind HFO propagation. Initially, we constructed co-activation matrices for each patient, derived from all recorded HFO sequences. Utilizing these matrices, we computed node hubness indicators, such as degree, betweenness, and eigenvector centrality, to evaluate the relative prominence within the SOZ nodes compared to those outside. Subsequently, we employed coherence analysis for each identified HFO sequence to quantify neural synchrony within the high-frequency band (>80 Hz), both within and surrounding the HFO-involved regions. These regions include the clinically defined SOZ by epileptologists and the HFO-zone (HFOZ), identified by the channels exhibiting HFO activity.

In order to assess the directional connectivity of high-frequency information, we applied the directed transfer function (DTF) analysis, grounded in multivariate autoregressive (MVAR) models *(26–30)*. DTF analysis provides a spectral perspective on the directionality of information flow in a multivariate system, facilitating the identification of pathological networks characteristic of epilepsy *(26, 31)*. This analysis was performed based on the clustered HFO sequences focusing on the information transmission in the high-frequency band (>80 Hz) for each patient, considering the consistency of the spatiotemporal patterns in the multi-channel events, using the Fieldtrip open-source toolbox *(32)*. The directional information analysis enabled a comparative examination of connectivity patterns across the HFOZ, SOZ, and peripheral regions, highlighting similarities and differences in information flow during HFO sequences. In addition, we identified a control group of non-propagating HFOs that, despite detection, did not exhibit consistent propagation patterns. These instances, characterized by localized high-frequency activity, provided a baseline for comparing connectivity strengths between propagating and non-propagating HFOs. For these activities, the local foci were defined by the extent of 15 mm from the HFOs channel. The directional information flow was quantified across each electrode in the intracranial EEG recording and between channel pairs within the high-frequency band. The channels were then grouped by regions of interest to form regional connectivity pairs for detailed analysis. To quantify the net directional information flow between the foci regions of interest and the peripheral areas, we employed the information asymmetry index, which is defined as the disparity in information exchange, normalized by the aggregate connectivity *(26, 27)*. This index quantifies the balance between incoming and outgoing information, with a positive index indicating a predominant outflow whereas a negative value denoting a prevalent inflow.

Further, the HFO sequences were segmented into initiation (early 33%) and progression (subsequent 67%) stages, and the peripheral regions were classified based on proximity (close: within ~15mm, intermediate: ~55mm, and distant: ~100mm). This categorization allowed for a detailed examination of information flow dynamics on a finer spatial scale, aiming to reveal the neural mechanisms governing the onset and progression of HFOs. We calculated the information flow exchanges between the initiation nodes and the progression nodes as well as the peripheral regions categorized as close, intermediate, and distant from the HFO-implicated areas, capturing both incoming and outgoing information of each node pair during the sequences. Additionally, the information asymmetry index was applied to measure the net directional information flow between these region pairs, elucidating the dynamic connectivity patterns that govern the propagation of HFOs *(26, 27)*.

### Figures

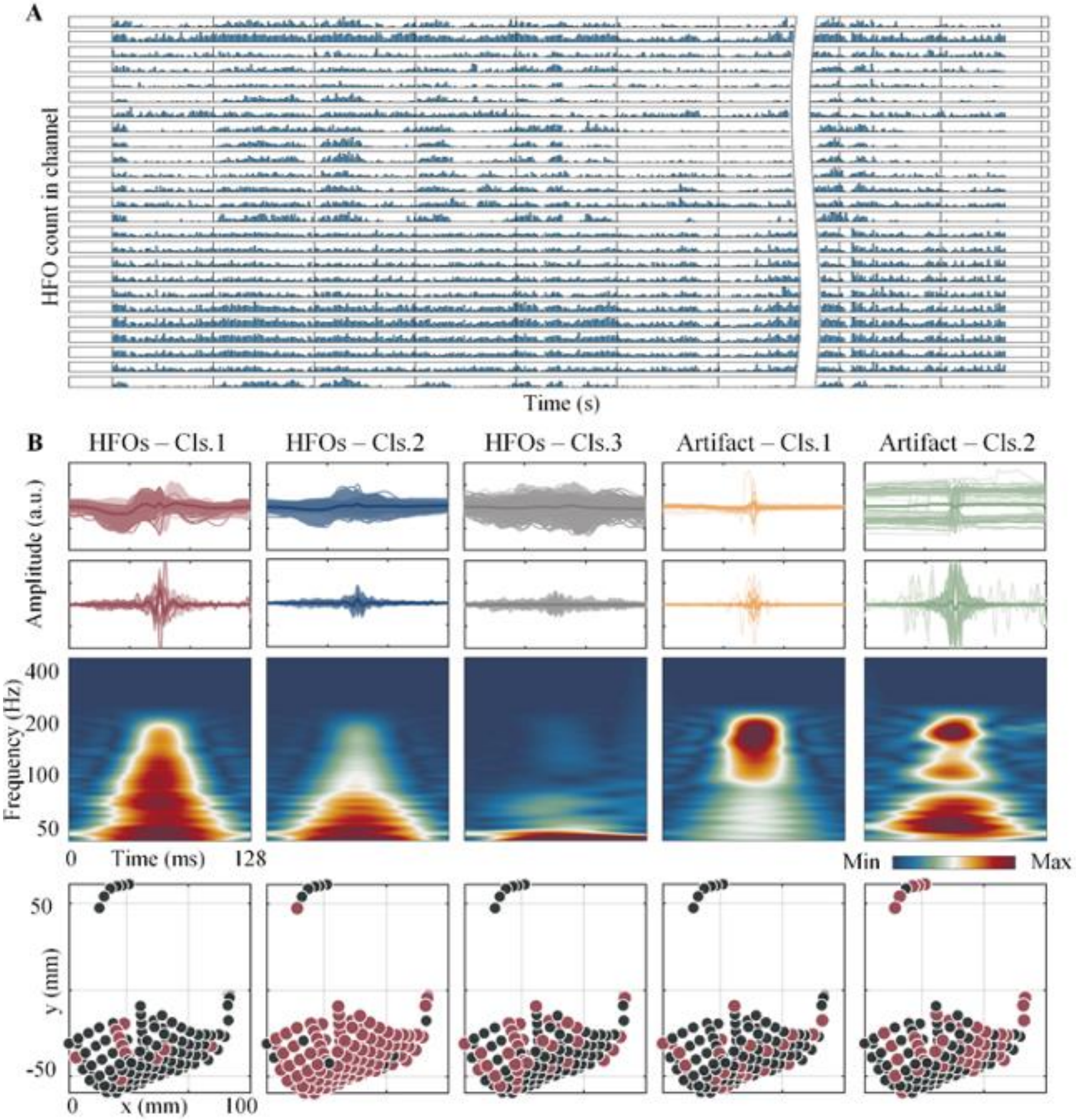


**Figure S1. Detection of HFOs in long-term intracranial EEG recordings. (A)** Histogram of detected HFOs across about a full-day long recording from a representative patient. **(B)** Clustered HFOs in each group are displayed with the raw data, high-pass filtered (>80 Hz) data, time-frequency representations, and spatial distribution over the iEEG electrodes located in a two-dimensional space modeled from post-implanatation CT. The signal traces are piled with the mean in bold and colors correspond to each group. Note that the first three clusters show clear HFOs in the filtered signals (lower panels) and unique low-frequency profile in the raw signals (upper panels), with the first cluster exhibiting potential interictal epileptiform discharges in the raw signal, and the last two clusters showing obvious groups of artifacts.

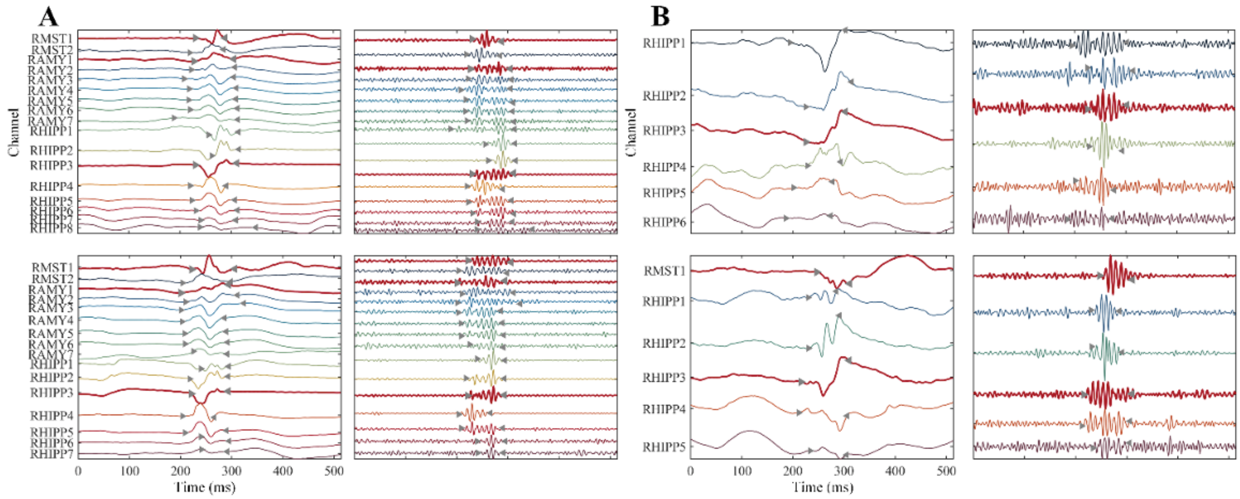


**Figure S2. Examples of identified HFO sequences in one sample patient. (A)** Two example HFO sequences with large extent, associated with cortical regions encompassing the amygdala, hippocampus, and middle temporal region. **(B)** Two HFO sequences with localized margin, involving mainly the hippocampus region. Note that within each group, (A and B), the HFO sequences (upper and lower panel) exhibit similar morphological characteristics, indicating a consistent pattern of neural activity; however, when comparing across groups (left and right panel), while there is a high spatial overlap in the involved electrodes (both recruiting the right hippocampus), the morphology and sequential order of the HFOs vary, underscoring distinct neural dynamics between these cortical areas. Bold red lines mark the SOZ channels, and left/right triangle indicate the onset/offset of the HFOs events.

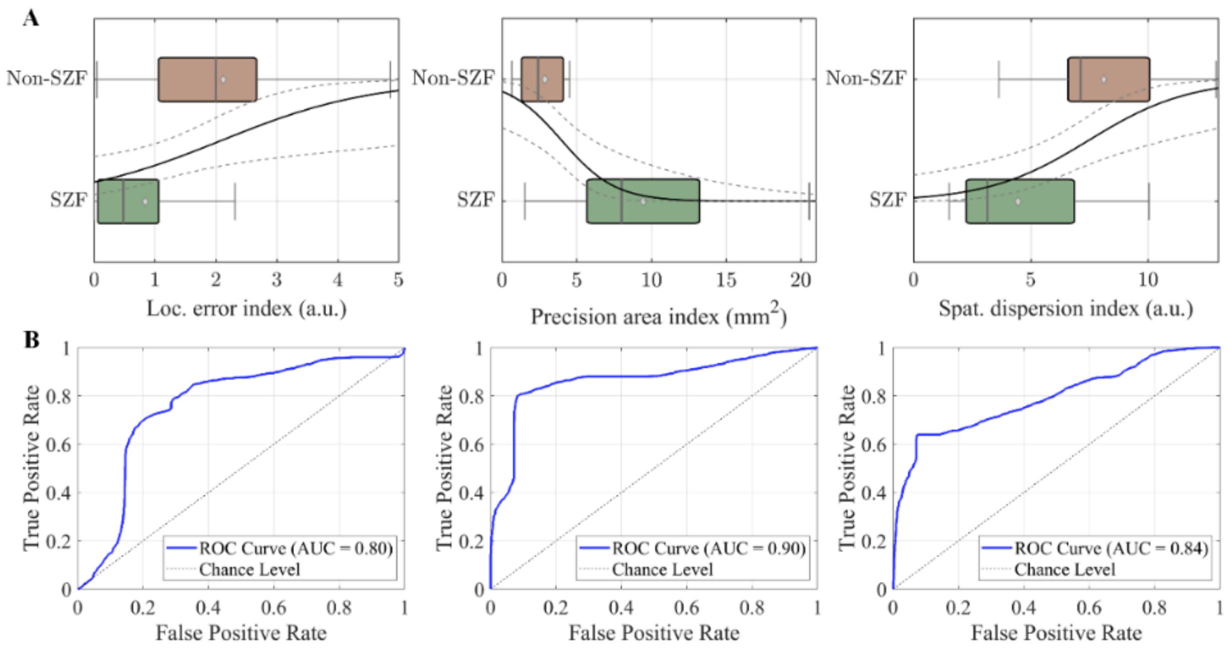


**Figure S3. Predictive analysis of seizure outcomes via HFO sequence mapping using generalized linear models. (A)** Association of estimated EZ of HFO sequences with post-surgical seizure outcomes. The relationship between localization measures (localization error, overlap ratio, and spatial dispersion) and surgical outcomes (seizure-free, SZF, and non-seizure-free, non-SZF) was modeled using generalized linear model in 39 patients (25 seizure-freedom). Box plots denote the median (dark line), mean (light grey diamond), interquartile range (IQR, horizontal bar), and range within 1.5 IQR (whiskers). The mean predictions are shown by solid dark curves with 95% confidence intervals (CIs) depicted by dashed curves. Displayed predictors include localization error (left), overlap ratio precision (center), and spatial dispersion (right), each normalized to the unit spatial resolution of intracranial electrodes. All metrics were statistically significant (all $p < 0.01$, Chi-squared test) in forecasting outcomes, with odds ratios of 2.22 (95% CI: 1.16–4.24) for localization error, 0.57 (95% CI: 0.39–0.85) for overlap precision, and 1.59 (95% CI: 1.17–2.18) for spatial dispersion indices per unit increase, predicting non-seizure freedom. **(B)** Receiver operating characteristic (ROC) analysis for the three aforementioned predictors. ROC curves are in blue, and the diagonal dashed line indicates chance performance, with the AUC (area under the curve) values denoted. The most effective model, using the overlap ratio precision index, yielded an AUC of 0.9, with accuracy, precision, recall, and specificity rates of 0.84, 0.95, 0.81, and 0.91, respectively. Note that all ROC curves strongly surpass the chance level, demonstrating predictive validity.

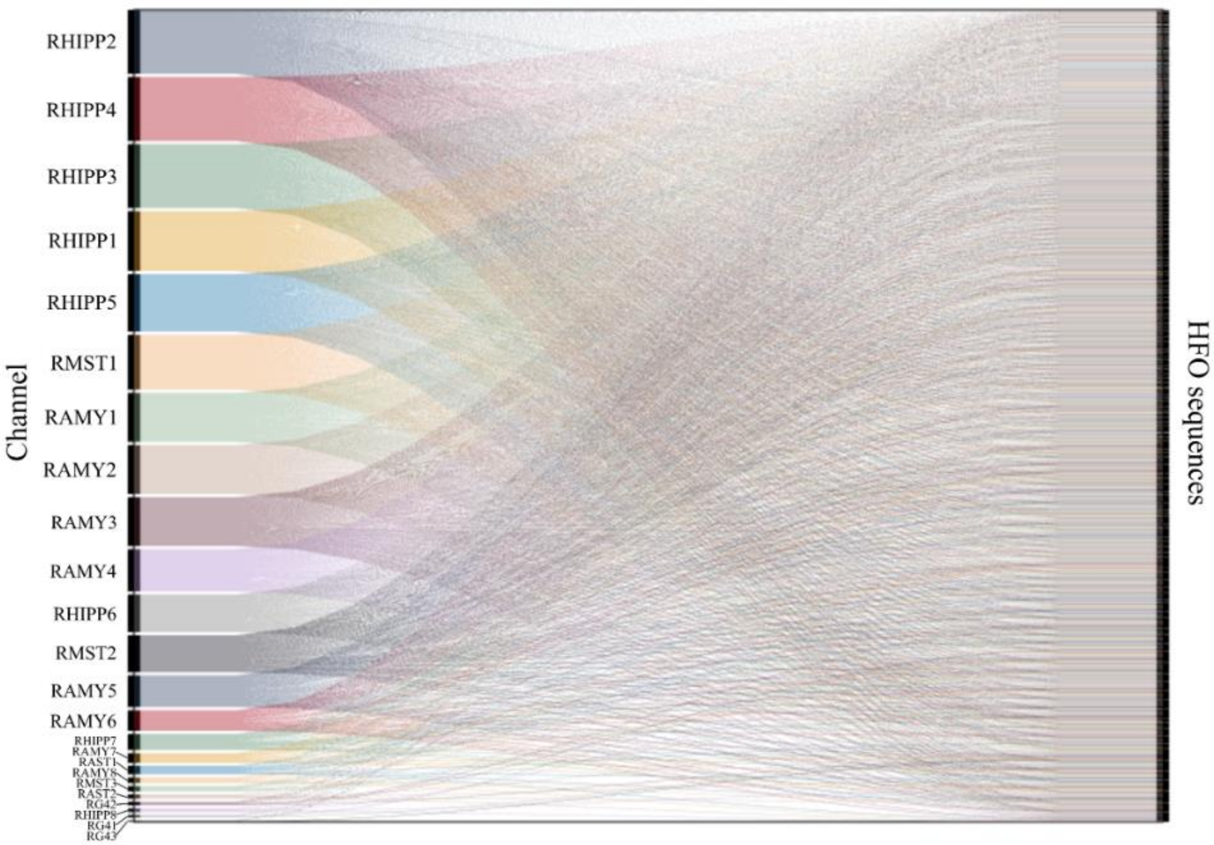


**Figure S4. Channel engagement in HFOs sequences.** Across the sequences (right vertical axis), multiple channels are repeatedly involved (left vertical axis), highlighting the consistent and predominant recruitment of the mesial temporal regions. Conversely, the lesser engagement of other channels (last few channels), which appear in only a minimal number of sequences, indicates sporadic recruitment in the associated cortical areas. RHIPP: right hippocampus, RAMY: right amygdala, RMST: right middle subtemporal, RAST: right anterior subtemporal , RG: right frontotemporal neocortex.

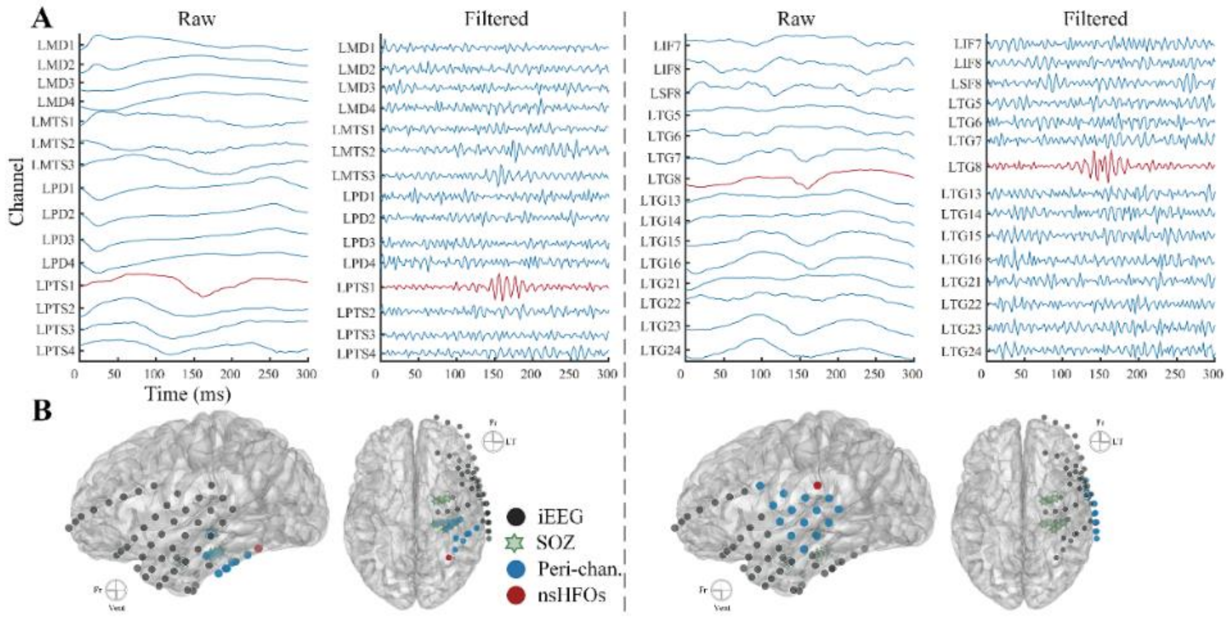


**Figure S5. Examples of non-propagating HFOs. (A)** Raw and high-pass filtered (above 80 Hz) EEG signals of non-propagating HFOs (red) and the activities at adjacent channels (blue, within ~40mm extent from the HFOs channel). **(B)** The spatial layout of intracranial EEG electrodes (dark), the channel with detected HFOs (red) and the surrounding channels (blue) as displayed in (A), and the SOZ marked with green hexagons on the cortical surface.
